# Phylotranscriptomics reveals discordance in the phylogeny of Hawaiian *Drosophila* and *Scaptomyza* (Diptera: Drosophilidae)

**DOI:** 10.1101/2021.07.08.451653

**Authors:** Samuel H. Church, Cassandra G. Extavour

## Abstract

Island radiations present natural laboratories for studying the evolutionary process. The Hawaiian Drosophilidae are one such radiation, with nearly 600 described species and substantial morphological and ecological diversification. These species are largely divided into a few major clades, but the relationship between these clades remains uncertain. Here we present 12 new assembled transcriptomes from across these clades, and use these transcriptomes to resolve the base of the evolutionary radiation. We recover a new hypothesis for the relationship between clades, and demonstrate its support over previously published hypotheses. We then use the evolutionary radiation to explore dynamics of concordance in phylogenetic support, by analyzing the gene and site concordance factors for every possible topological combination of major groups. We show that high bootstrap values mask low evolutionary concordance, and we demonstrate that the most likely topology is distinct from the topology with the highest support across gene trees and from the topology with highest support across sites. We then combine all previously published genetic data for the group to estimate a time-calibrated tree for over 300 species of drosophilids. Finally, we digitize dozens of published Hawaiian Drosophilidae descriptions, and use this to pinpoint probable evolutionary shifts in reproductive ecology as well as body, wing, and egg size. We show that by examining the entire landscape of tree and trait space, we can gain a more complete understanding of how evolutionary dynamics play out across an island radiation.

## Introduction

In the era of genome-scale data, we have an opportunity to unpack the biological meaning of phylogenetic support. In phylogenetic analyses that seek to discover the relationships between organisms, support is often defined as the proportion of information that favors a particular branch in an evolutionary tree^1^. Methods have been developed that emphasize extracting the tree with the greatest amount of support from out of an otherwise rugged landscape of treespace^2,3^. However, a growing number of studies have emphasized the biological relevance of that landscape to our understanding of the evolutionary process^4–6^. For example, many new studies have contributed evidence that, even with trees with high measures of conventional support, we can expect large amounts of discordance among sites and genes, especially when examining speciation events with short internodes or with a likelihood of introgression^7,8^. Here we use the island radiation of Hawaiian drosophilid flies to study the landscape of treespace, and show that the relationships between the major groups of these flies are best understood by using methods that embrace evolutionary discordance.

The Hawaiian *Drosophila* have a long history as a model clade for the implementation of phylogenetic methods^9^. More than twenty years ago, Baker and Desalle used the Hawaiian radiation of *Drosophila* to perform one of the first analyses to demonstrate incongruence between an overall species tree and underlying gene trees^10^. Their study focused on the resolution between major clades of Hawaiian *Drosophila* and built on the landmark work done by Carson in the 1970s inferring the phylogeny of a subgroup of Hawaiian *Drosophila*, the picture-wing flies, based on the banding pattern of polytene chromosomes^11^, among other early phylogenetic studies^12,13^. During the past twenty years, the relationships between major groups has been revisited several times^14,15^. Most recently, O’Grady and colleagues (2011) used mitochondrial genes and expanded taxon sampling^16^, and Magnacca and Price (2015) used an expanded nuclear gene set^17^. The study presented here builds on this foundational work, presenting the first phylogenetic analysis of genome-scale data for the group.

The Hawaiian Drosophilidae consist of 566 described species^18,19^, with hundreds more estimated to be awaiting description^18^. These species have been divided into the following major clades^18^: [1] the *picture-wing, nudidrosophila, ateledrosophila* (PNA) clade, which has served as a model clade for the study of sexual selection^20^ and speciation^21^; [2] the *antopocerus, modified-tarsus, ciliated-tarsus* (AMC) clade, first proposed by Heed (1968), ^18,22^and confirmed by subsequent phylogenetic studies^16,23^; [3] the *modified-mouthparts* (MM) clade; and [4] the *haleakalae* clade, an enigmatic group in need of further study^24^. Several other smaller clades have been suggested as falling outside of these major groups, including the *rustica* group of three species^25^, and the monotypic lineages of *D. primaeva* and *D. adventitia*. The position of *D. primaeva* has been somewhat uncertain, but several studies have suggested it is the sister taxon to *picture-wing* flies^14^, including the work on polytene chromosomes by Carson and Stalker^26^. The species *D. adventitia* was originally suggested to be part of the MM clade^27^, but recent studies placed it as the sister taxon to *D. primaeva*^14^or possibly other major clades. Additionally, the Hawaiian *Drosophila* are the sister clade of the genus *Scaptomyza*, which is nested within the broader paraphyletic genus *Drosophila* and is hypothesized to have colonized the island independently^28,29^, possibly more than once^30^. Throughout this manuscript, we use Hawaiian *Drosophila* to refer to non-*Scaptomyza* Hawaiian species, and Hawaiian Drosophilidae to refer to the clade of Hawaiian *Drosophila*+*Scaptomyza*.

Many phylogenetic studies have been performed which have confirmed the monophyly of each of these clades and provided resolution for internal relationships (PNA^17,31^, AMC^23,32^, *haleakalae*^33^, and *Scaptomyza*^29,30^). Previous phylogenetic studies, however, have not resulted in a consensus relationship between the major clades within Hawaiian *Drosophila* (Fig. S1)^17^. Magnacca and Price (2015) showed that different phylogenetic methods of analysis (e.g., using software based on Bayesian statistics rather than maximum likelihood for inference) produced highly incongruent topologies (Fig. 1)^17^. In that study, the most likely topology had *D. primaeva* as the sister taxon to all other Hawaiian *Drosophila*, and included a clade uniting MM+AMC+*haleakalae*, with the *haleakalae* clade showing greater affinity to AMC species relative to MM species (Fig. 1B). This topology was consistent with the tree suggested by O’Grady and colleagues (2011) analysing mitochondrial data and using maximum likelihood and Bayesian analyses^16^. However the analyses of Magnacca and Price (2015) using Bayesian software package BEAST showed an alternative relationship, with *haleakalae* flies as the sister clade to all other Hawaiian *Drosophila*, a clade uniting the MM+PNA+*D. primaeva*, and an closer affinity between *D. primaeva* and PNA species than between *D. primaeva* and MM species (Fig. 1C). This latter arrangement is largely consistent with relationships proposed by Throckmorton in 1966^28^ and reiterated in several subsequent studies (Fig. S1)^10,14,15^.

**Figure 1:**
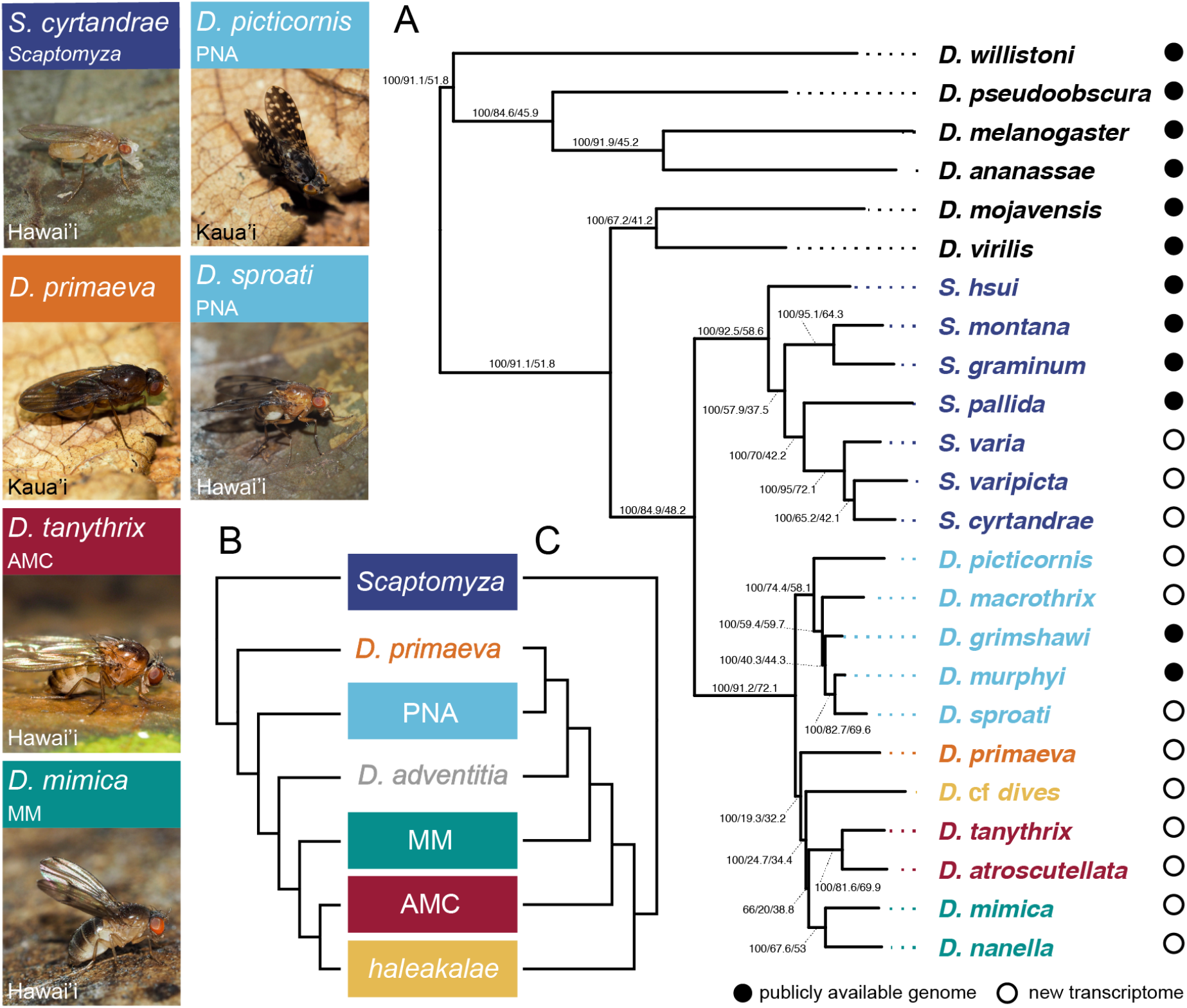
Phylotranscriptomic analysis indicates relationships between major clades distinct from those previously hypothesized. Photos show six of the twelve species with *de novo* transcriptomes presented in this study, listing their parent clade and the Hawaiian island on which they are found. A, Results novel to this study, showing best supported tree across maximum likelihood and Bayesian analyses. Node labels indicate ultrafast bootstrap values / gene tree concordance factors (gCF) / site concordance factors (sCF), see concordance factor analysis below. *D. adventitia* was not present in phylotranscriptomic analyses, see Fig. 3 for information on its placement. B-C, Previously hypothesized relationships between the *picture wing*-*nudidrosophila*-*ateledrosophila* (PNA), *modified-mouthparts* (MM), *antopocerus*-*modified tarsus*-*ciliated tarsus* (AMC), *haleakalae*, and *Scaptomyza* clades, as well as two monotypic clades, *D. primaeva* and *D. adventitia*. Topology B was recovered in O’Grady and colleagues (2011) ^16^and Magnacca and Price (2015)^17^. Topology C was recovered using the Bayesian software BEAST in Magnacca and Price (2015)^17^, showing incongruent relationships between clades at the base of the radiation of Hawaiian *Drosophila*.

Resolving these relationships is critical for our understanding of the morphological and ecological evolution of these flies^14–16^. Hawaiian *Drosophila* demonstrate a large diversity in body size^34^, wing size^35^, and egg size^36^; in the number and position of structural features such as wing spots^35^; in the number of egg-producing units in the ovary (ovarioles)^37,38^; and in the type of substrate used for oviposition and larval feeding^15,39^. Some clades demonstrate unique suites of morphological and behavioral traits, whose evolutionary history is unclear because of uncertainties in the phylogeny. For example, the *haleakalae* flies exclusively use fungal oviposition substrates and are considered to have less complex mating behaviors than other, more well-studied groups (e.g., *picture-wing* flies)^24^. It is unclear whether this suite of traits represents a secondary transition relative to the ancestral state, because it is not known whether *haleakalae* flies are the sister clade to all other Hawaiian *Drosophila* or nested within the radiation. Resolution in the relationships at the base of this lineage will be key in identifying which branches experienced substantial trait diversification, and especially in identifying whether any of these traits demonstrate predictable patterns of co-evolution.

Here we present the first phylogenomic relationships between the major groups of Hawaiian Drosophilidae. We combine twelve new transcriptomes sequenced in this study with recently published genomes for two Hawaiian *Drosophila* species^40^, four non-Hawaiian *Scaptomyza*^40^, and six outgroup species^41^. By increasing the number of genes used to infer relationships, we begin to unpack the evolutionary history in the short internodes at the base of the Hawaiian *Drosophila* radiation. Following up on the critical study by Baker and Desalle 25 years ago^10^, we explore the landscape of treespace and the discordance between species and gene trees using our phylotranscriptomic dataset. We then use the results of our analysis as initial constraints on subsequent phylogenetic analyses using a dataset of 316 species and 44 genes, compiled using all previous phylogenetic studies of Hawaiian Drosophilidae. Finally, we estimate the age of the radiation, and use this time-calibrated tree to identify branches where shifts in trait evolution likely occurred. Our findings suggest a relationship between major clades that is distinct from both previously hypothesized topologies, and that is well supported by both maximum likelihood and Bayesian analyses. We show that examining a comprehensive landscape of tree and trait space can allow for a more complete understanding of evolutionary dynamics in this remarkable island radiation.

## Results

### Phylotranscriptomics suggest a new phylogeny of Hawaiian Drosophilidae

Using a phylotranscriptomic approach, we recovered a new topology between the major clades of Hawaiian Drosophilidae, distinct from those previously hypothesized (Fig. 1, S1). This topology was the most likely tree estimated using IQtree^42^ and RAxML^43^, as well as the consensus tree with highest posterior probability estimated using PhyloBayes^44^ (Fig. 1A, S2, S3). Bootstrap support for all branches was 100 and posterior probability was 1, with the exception of the branch subtending the clade uniting MM+AMC (IQtree ultrafast bootstrap: 66, RAxML bootstrap: 57, PhyloBayes posterior probability: 0.52). We also estimated the phylogeny using a multi-species coalescent model with ASTRAL^45^, and recovered the same topology with the exception of the placement of *D. primaeva* (as the sister taxon to PNA, Fig. S4). Each of these analyses were performed on a supermatrix of 10,949 putatively orthologous genes, aligned and assembled using the agalma pipeline^46^ with no filtering based on occupancy (actual gene occupancy was 41.7%). To test the senstivity of our results to missing data, we repeated the IQtree analysis on a dataset reduced using an occupancy threshold that ensures representation of 80% of taxa at each gene (1,926 genes), and recovered the same topology as with the full set of genes (Fig. S5).

The most likely tree indicates that the PNA clade, including *picture-wing* species, is the sister clade to all other Hawaiian *Drosophila. D. primaeva* is found to be the sister taxon to a clade containing non-PNA Hawaiian *Drosophila*, though this clade received lower support when using the dataset reduced by occupancy (Fig. S5, ultrafast bootstrap of 85). A second monotypic lineage, *D. adventitia*, was not sampled for phylotranscriptomic analyses, but using specific gene markers, we recover this as the sister taxon to a clade including AMC+MM+*haleakalae* (see section on expanded phylogenetic analysis below). This latter clade was previously recovered in previous phylogenetic analyses^16,17^. In contrast to those studies, which suggested a monophyletic clade of AMC+*haleakalae*, we do not recover sufficient support for any particular arrangement of MM, AMC, and *haleakalae* (ultrafast bootstrap from both the full and reduced occupancy matrix is <95).

We tested the most likely tree emerging from our analysis (Fig. 1A) against two previously suggested alternative hypotheses (Fig. 1B-C) using the Swofford-Olsen-Waddell-Hillis (SOWH) test^2^, a parametric bootstrap approach for comparing phylogenetic hypotheses. In both cases, the difference in likelihood between the most likely tree and these alternatives was larger than we would expect by chance (p-value for both <0.01, with a sample size of 100). Between Fig. 1A and 1B the difference in log-likelihood was 1774.1, and between Fig. 1A and 1C was 6132.1, while the null distribution according to the SOWH test had no differences greater than 15 for either comparison. Taken together, our results suggest a new phylogeny for Hawaiian Drosophilidae relationships wherein MM, AMC, and *haleakalae* represent a monophyletic group, and the PNA clade, rather than either the *haleakalae* clade or *D. primaeva*, is the sister clade to all others (Fig. 1A).

### Identifying hotspots of gene and site concordance in treespace

We analyzed the strength of phylogenetic concordance in our phylotranscriptomic dataset by estimating the gene and site concordance factors for each branch in our tree. Gene concordance factors (gCF) are calculated as the proportion of informative gene trees that contain a given branch between taxa, and can range from 0 to 100^5,47^. Site concordance factors (sCF) are calculated as the average proportion of informative sites that support a given branch between taxa. Because one site can only support one of three arrangements for a quartet of taxa, sCF typically ranges from ∼33.3 to 100, with 33.3 representing our null expectation based on chance^47^. We found that for many branches in our tree both gCF and sCF are high, indicating these relationships are supported by a majority of gene and sites in our dataset. For example, the branch uniting Hawaiian *Drosophila* has a gCF of 91.2, and sCF of 72.1 (Fig. 1A). However for the branches subtending most relationships between the major clades of Hawaiian *Drosophila*, gCF and sCF are low. For example, the branch uniting *D. primaeva*+*haleakalae*+AMC+MM to the exclusion of PNA has a bootstrap value of 100, but a gCF of 19.3 and sCF of 32.2.

We also tested the extent to which potential error in multiple sequence alingment affected concordance values by filtering out poorly aligned sequence fragments, and repeating the tree inference and concordance analyses. After filtering poorly aligned sequences, we recover the same topology as Fig 1A, with the exception of the arrangement of MM, AMC, and *haleakalae*, here showing a MM+*haleakalae* as monophyletic (IQtree ultrafast bootstrap: 97, Fig. S6). Concordance factors between analyses on filtered and non-filtered data were nearly equivalent (e.g. the branch separating PNA from the other Hawaiian *Drosophila* received a gCF of 19.0 and sCF of 31.6 after filtering, and a gCF of 19.3 and sCF of 32.2 using all data). Furthermore, using a series of stringency thresholds to filter the data, we observed no pattern of increasing or decreasing concordance factors across branches (Fig. S7). These results suggest that values of discordance in this phylogeny are not artificially inflated due to technical errors from the alignment step.

We interpret the measures of discordance as reflecting real variation in the phylogenetic signal of different genes and sites, which is not unexpected for a radiation such as this with short internodes subtending major clades^47^. Furthermore, the presence of discordance does not mean that there is little that can be said about the relationships between these groups. In contrast, by unpacking this discordance we can begin to qualitatively describe the amount and distribution of phylogenetic signal for multiple alternative, plausible bipartitions.

To this end, we first visualized hotspots of concordance across treespace (Fig. 2). We created all 105 topological combinations of the possible arrangements between major clades, and then re-estimated gCF and sCF for each. Visualizing the mean values for gCF and sCF plotted in treespace shows that the most likely tree, as estimated with IQtree, is not the tree with the highest mean gCF and sCF, but it is near a hotspot of alternative arrangements for which both of these values are high (Fig 2, treespace, most likely tree indicated by dark red outline). In contrast to the most likely topology, the trees with the top three mean gCF values and two of the three trees with the top mean sCF values unite *D. primaeva*+PNA to the exclusion of other Hawaiian *Drosophila*. Variation between these top trees largely depends on the placement of *haleakalae* relative to other clades (Fig. 2, top gCF and sCF trees).

**Figure 2:**
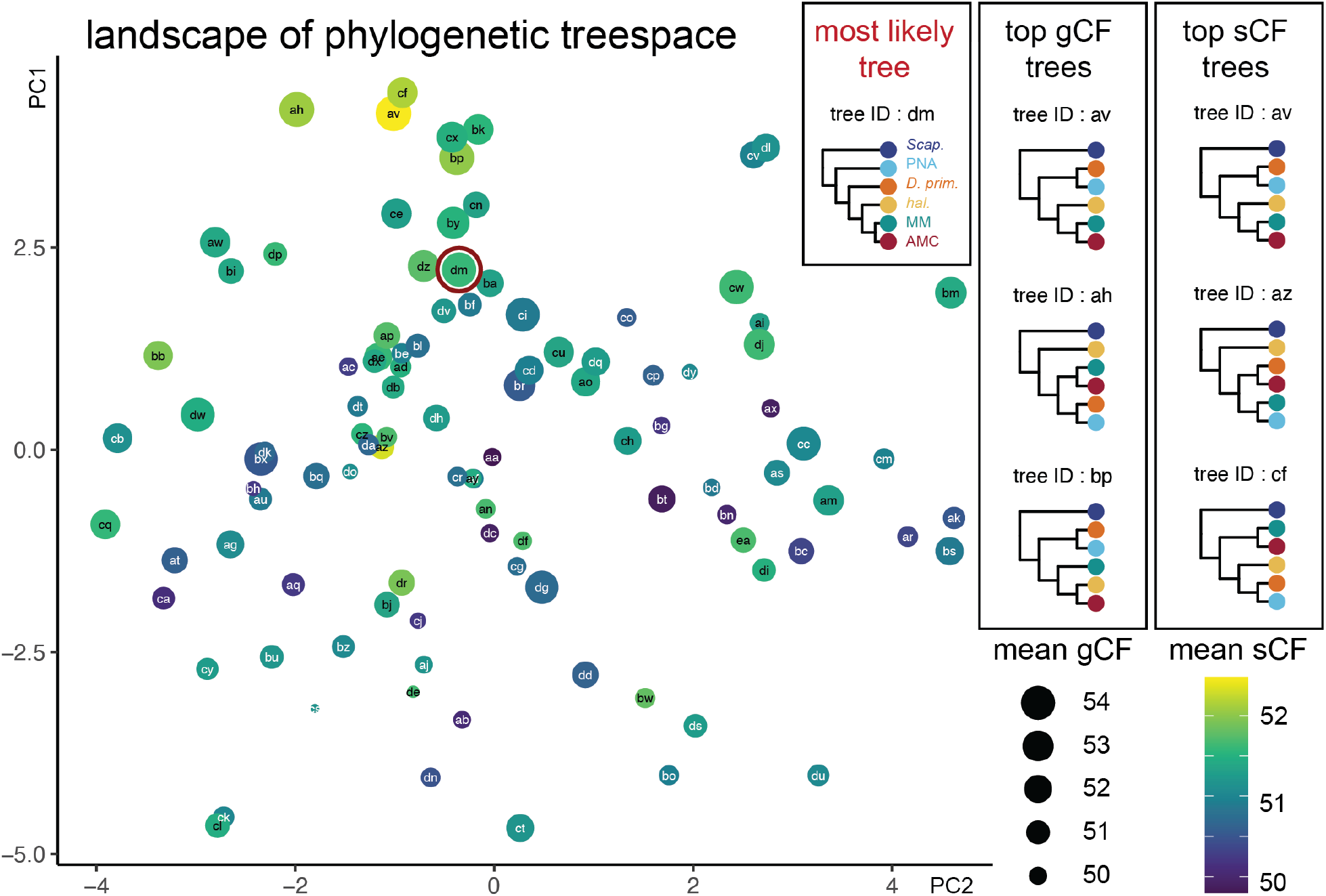
The landscape of treespace shows hotpots of concordance among genes and sites. The landscape of treespace for all possible topological combinations of the five clades of Hawaiian *Drosophila* studied here: PNA, *D. primaeva, haleakalae*, MM, and AMC. Individuals points represent different arrangements of the five clades, labeled randomly with two-letter IDs from aa through ea. The distance between points indicates tree similarity (calculated from Robinson-Foulds distances). The size of points represents the mean gene concordance factor (gCF) across relevant branches, and the color represents the mean site concordance factor (sCF, purple=low, yellow=high). The point outlined in red (tree dm) indicates the best topology found with IQtree, RAxML, and PhyloBayes, which is distinct from the top trees according to mean gCF (av, ah, and, ap) or mean sCF (av, az, and cf). Concordance measurements for all topologies are available, see Methods and Data availability.

The mean gCF and sCF across branches may not always be informative metrics, given that some topologies may contain one highly supported branch and others with very low support. Therefore, we also analyzed concordance for all the unique bipartitions across the set of possible topologies (Figs. S8 and S9, see Supplemental Section ‘Concordance Factor Analysis’). We found that for gCF, there is clear signal supporting bipartitions that unite *D. primaeva*+PNA, as well as those that unite MM+AMC+*haleakalae* (Fig. S8). We found that for sCF, concordance values across bipartitions are more variable, but those that unite PNA+*haleakalae* show less support than we might expect by chance, while those that unite *D. primaeva*+PNA and AMC+MM show more support (Fig. S9). In addition, between gCF and sCF, we found conflicting signals for bipartitions that define one clade as sister to the rest of Hawaiian *Drosophila*, with gCF indicating support for PNA (consistent with the most likely topology), and sCF indicating support for *haleakalae*.

To investigate the source of discordance across genes, we performed a likelihood mapping analysis that assesses the phylogenetic information in each gene^42,48^. This analysis calculates the likelihood support for the three possible arrangements of each quartet of taxa in an alignment, and then counts the number of informative quartets that strongly support one arrangement over the other two^48^. Here we performed likelihood mapping on each of the 10949 genes in our full dataset, resulting in 2182 quartets relevant to the position of the PNA group and 3075 quartets relevant to the position of the *haleakalae* group. Our results showed that, in both cases, the vast majority of quartets are uninformative, with no strong likelihood support for any one arrangement (Fig. S10). While support was for the most part evenly divided among possible arrangements, we observed more quartets uniting *haleakalae*+MM, to the exclusion of AMC, PNA, and *D. primaeva*, as well more quartets uniting PNA+*D. primaeva* to the exclusion of other Hawaiian Drosophilidae (Fig. S10A,C). We also tested whether genes that support one topology over another at these nodes were enriched for either long or short genes, or fast or slow-evolving genes. Our results showed no correlation between gene tree topology and gene length or evolutionary conservation (Fig. S11), with the exception of genes supporting *D. primaeva* as the sister to all other Hawaiian *Drosophila*, which was supported by somewhat slower evolving genes than alternative arrangements (Fig. S11D).

In summary, across all analyses we found consistent evidence for a bipartition that separates PNA from clades that include MM and AMC. While the placement of *D. primaeva* was strongly supported in our maximum likelihood and Bayesian analyses, we observed substantial discordance in this arrangement, and detect signal suggesting a significant amount of shared history between *D. primaeva* and PNA. Similarly, while the clade uniting MM+AMC+*haleakalae* received strong bootstrap support, we observed substantial discordance in the placement of *haleakalae*, and suggest that further resolution in its placement will be possible with additional taxon sampling in that clade.

### Calibrating an expanded phylogeny to time

Building on the phylotranscriptomic analyses above, we collected all publicly available genomic and transcriptomic data for species from Hawaiian *Drosophila* and *Scaptomyza*. These data were accessioned in nine analyses published since 1997, most of which focused on resolving the phylogenetic relationships within a major clade^10,17,23,29,29–33^. The dataset we compiled contained 44 genes (6 mitochondrial and 38 nuclear) from 316 species (including 271 described and 45 undescribed putative species), with an overall occupancy of 17.3% (Fig. S12). We used this dataset to infer the phylogeny with IQtree, constraining the relationships between major clades to conform to the topology shown in Figure 1A.

The resulting topology is to our knowledge the most species rich phylogenetic tree of the Hawaiian Drosophilidae to date (Fig. S13). Several support values are low (ultrafast bootstrap <95), especially for nodes near the base of the radiation. However, this is not unexpected, given that this phylogeny is estimated primarily from the same data previously analyzed, which recovered alternative relationships at those nodes. Of note are the low support values for the relationships within the MM and *haleakalae* clades (Fig. S13, polytomies), emphasizing the need for further study in these groups.

We used this expanded genetic dataset and topology to estimate the age of the Hawaiian Drosophilidae by calibrating this tree to time using the software package BEAST^49^. Consistent with recent publications^17,30,50^, our results indicate that the age of the split between Hawaiian *Drosophila* and *Scaptomyza* occurred between 20 and 25 million years ago (Fig. 3, median root age 22.8 million years). The results shown here were calibrated using updated estimates for the ages at which Hawaiian islands became habitable, based on models of island emergence, growth, and decline via erosion and subsidence^51^ (Table 1). Similar results were obtained using the calibration scheme from Russo and colleagues (2013) ^52^that includes a single fossil calibration point for the clade, based on the taxon *S. dominicana* recovered from dominican amber (median root age 22.9 million years). However, with both schemes uncertainty around the root age remains substantial (95% highest posterior density confidence interval 17.4 - 29 million years), and small changes in the calibration times used can lead to substantial differences in this estimate. When calibrating the tree using the same island age estimates as in Magnacca and Price (2015), which are marginally younger (Table 1), we estimated the age of Hawaiian Drosophilidae to be ∼15 million years old (median root age 15.5 million years). Furthermore, we note that calibrating using primarily vicariance based estimates of time is considered to be imprecise and should be avoided^53^. Taken together, we consider this estimate of the age of Hawaiian *Drosophila*, as well as those previously published, to be tentative, and suggest that further data (e.g., new fossil evidence) will be necessary to determine the age of diversification relative to island emergence with greater certainty.

**Figure 3:**
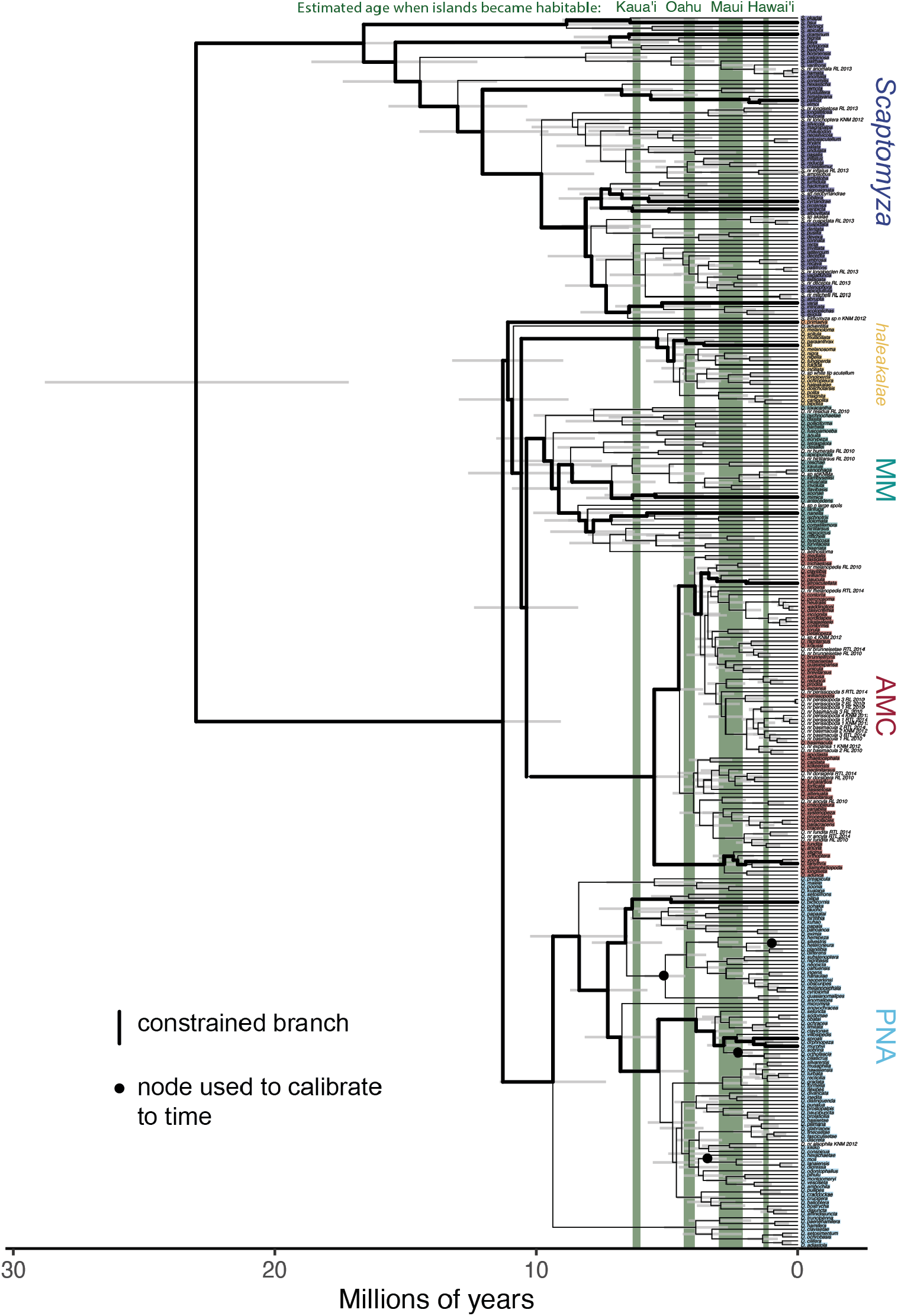
Time-calibrating the phylogeny of 316 Drosophilidae species. This phylogeny was inferred using IQtree to analyze all publicly available genetic data for Hawaiian *Drosophila* and *Scaptomyza*. It was then calibrated to time using the software BEAST, with four calibration points at nodes that show a biogeographic progression rule^17^. Similar results were obtained using a calibration scheme^52^ that takes into acount a single fossil taxon for the group (Table 1). The 95% highest posterior density intervals for each node are shown as gray bars, indicating the credible interval for the age of that group. The age at which four Hawaiian islands are estimated to have become habitable is shown in green. Colored labels indicate the clade to which taxa belong, and colors correspond to Fig 1; taxa without a colored label are species with genetic data that are as of yet undescribed. See Fig. S13 for bootstrap support. Calibration using only island biogeography is known to be imprecise^53^, therefore the divergence times shown here are considered tentative.

**Table 1:**
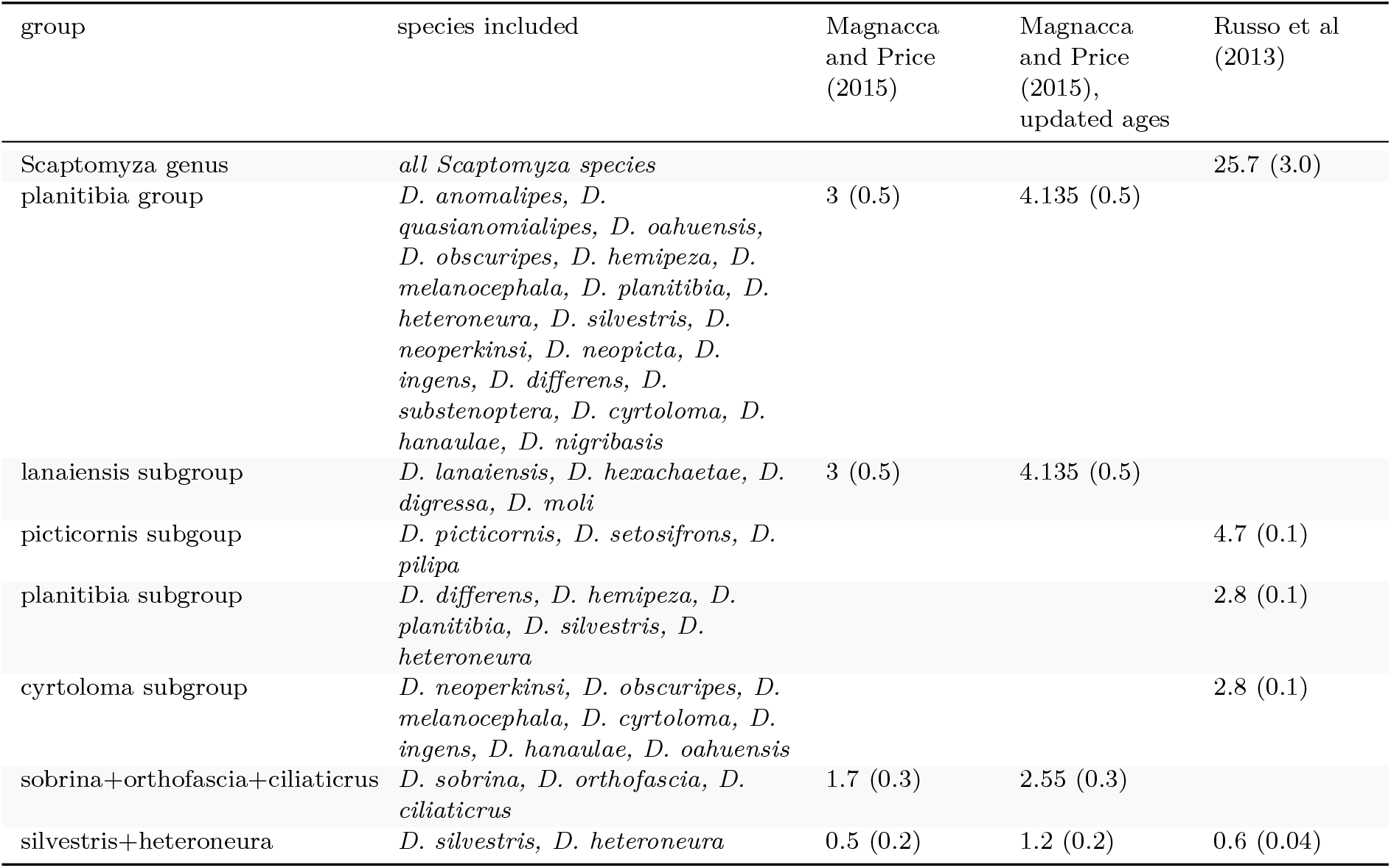
Calibration points for dating with BEAST. Values are mean (standard deviation) age in million years for normally distributed time priors.

According to this estimate, we find that the division between major Hawaiian *Drosophila* clades occurred around ten million years ago (Fig. 3), prior to the estimated time when the Hawaiian island Kaua’i became habitable (between 6.3 and 6.0 million years ago^51^). Our results show that the diversification of lineages within MM also occurred around that time, while the lineages within the AMC, *haleakalae*, and *grimshawi* groups (PNA) all arose within the last five million years, around the time Oahu became habitable. We note that the MM groups suffers from lower representation across genes used to calibrate the tree to time (Fig. S12), and suggest that more data may help shed light on differences in the age of this clade relative to others.

### Ancestral state reconstruction of oviposition and larval feeding ecology

With this time-calibrated tree for 316 species, we have an opportunity to investigate the evolutionary dynamics of trait diversification. By modeling the evolution of the diverse suite of ecological and morphological features across the phylogeny, we can identify which lineages have experienced major shifts in trait evolution. Predicting the number and phylogenetic position of these shifts will in turn be critical for informing future studies on development, life-history, and evolution of these flies. In the following analyses of trait evolution, we used the maximum clade credibility tree from the constrained BEAST analysis described above. Using this tree allows us to maximize the number of taxa for which genetic data are available, painting the most complete picture of ecological and morphological evolution in these flies up to this date. However, due to the fraction of genetic data missing across taxa, it also includes nodes with low bootstrap support (Fig. S13, polytomies). Therefore, for internal lineages for which evolutionary relationships remain uncertain, the position of these evolutionary shifts are subject to change as more genetic data become available and further phylogenetic resolution is achieved.

The Hawaiian Drosophilidae use a wide variety of plant, animal, and fungal species for egg laying and larval feeding (Fig. 4)^22,39,54^. The majority of species breed in rotting substrates, with variation in the part of the plant or fungus in question, including rotting bark, leaves, flowers, and fruit. A few species breed on live tissue, and one notable *Scaptomyza* subgenus, *Titanochaeta*, have been reared exclusively from spider egg masses^55^. In 2008, Magnacca and colleagues reviewed host plant and substrate records and found that, while many species can be considered specialists to species or substrate, host shifting was common and many species occasionally use non-preferred substrates^39^. The type of oviposition substrate has been suggested as a driver for diversification of the reproductive traits ovariole number and egg size^15,37,38^. However, the previous reconstruction of oviposition substrate by Kambysellis and colleagues (1997) ^15^ was performed with a phylogeny that included only three non-PNA species, and was therefore unable to resolve the ancestral oviposition substrate for Hawaiian *Drosophila* or to identify when evolutionary shifts in substrate outside of PNA were likely to have occurred.

**Figure 4:**
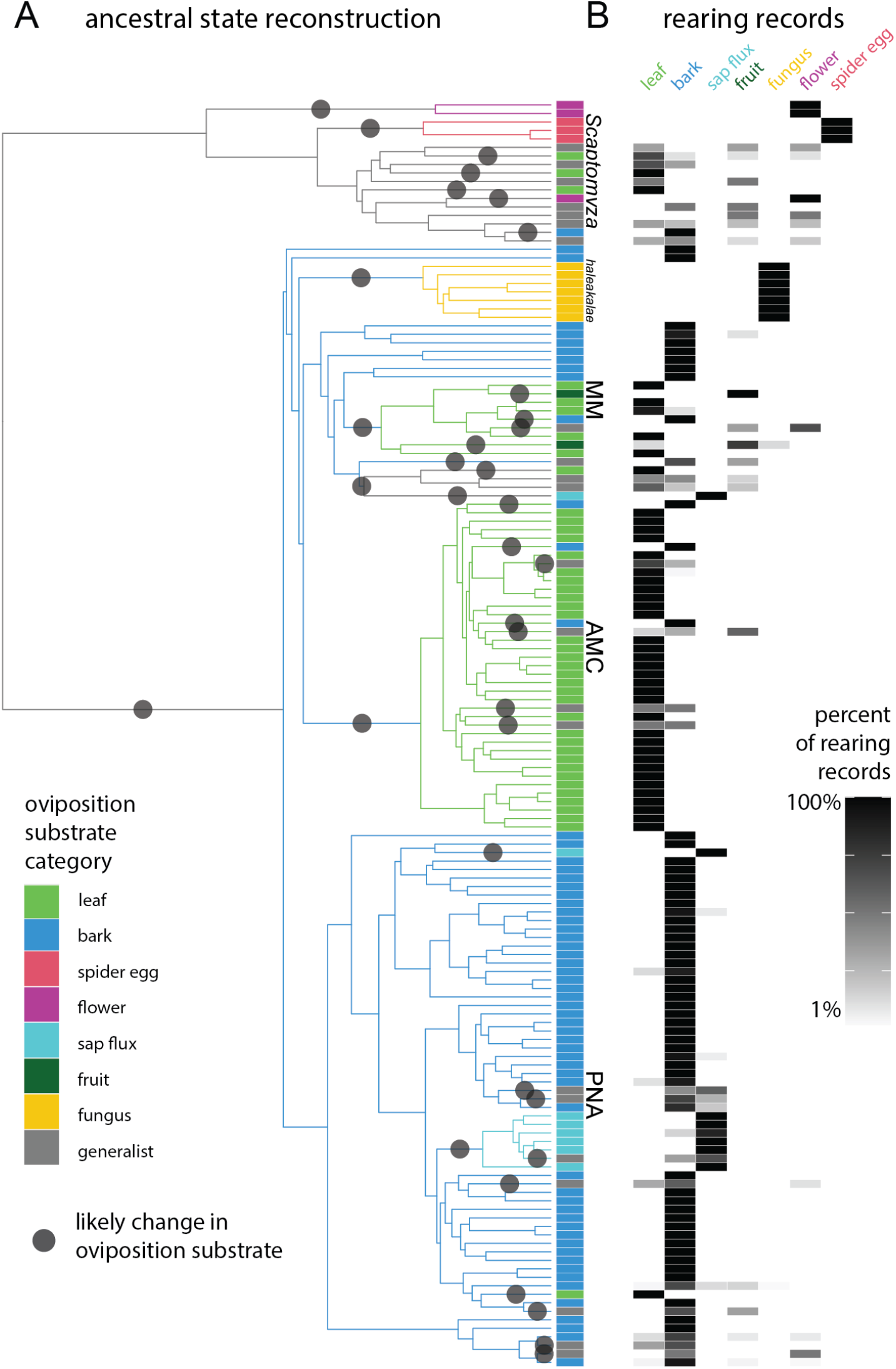
Ancestral state reconstruction of oviposition substrate indicates dozens of evolutionary transitions. A, We used stochastic character mapping to reconstruct the ancestral substrate used for oviposition and larval feeding, and identified dozens of likely transitions in substrate (gray circles). Branch color indicates the ancestral substrate type with highest probability, and tip box indicates extant oviposition substrate. B, Oviposition substrate category was defined based on rearing records, using the data summarized in Magnacca and colleagues (2008)^39^. Generalist species are defined as those with any two substrates that each comprise 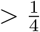 of rearing records, or any species with no substrate that comprises 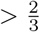 of rearing records.

We combined the phylogenetic results presented here with the data summarized in Magnacca and colleagues (2008), to reconstruct the ancestral oviposition substrate for the Hawaiian Drosophilidae (Fig. 5A, S14). Using stochastic character mapping^56^, we recover the most probable ancestral oviposition substrate for the Hawaiian *Drosophila* as bark breeding (defined as including rearing records from bark, stems, branches, roots, and fern rachises, see Supplemental Table S5). We recover a transition from bark to leaf breeding at the base of the AMC clade that has generally persisted throughout the diversification of that group. As previously reported^39^, we find several groups that demonstrate no reported variation in substrate type (e.g., fungus breeding *haleakalae*, Fig. 5B).

**Figure 5:**
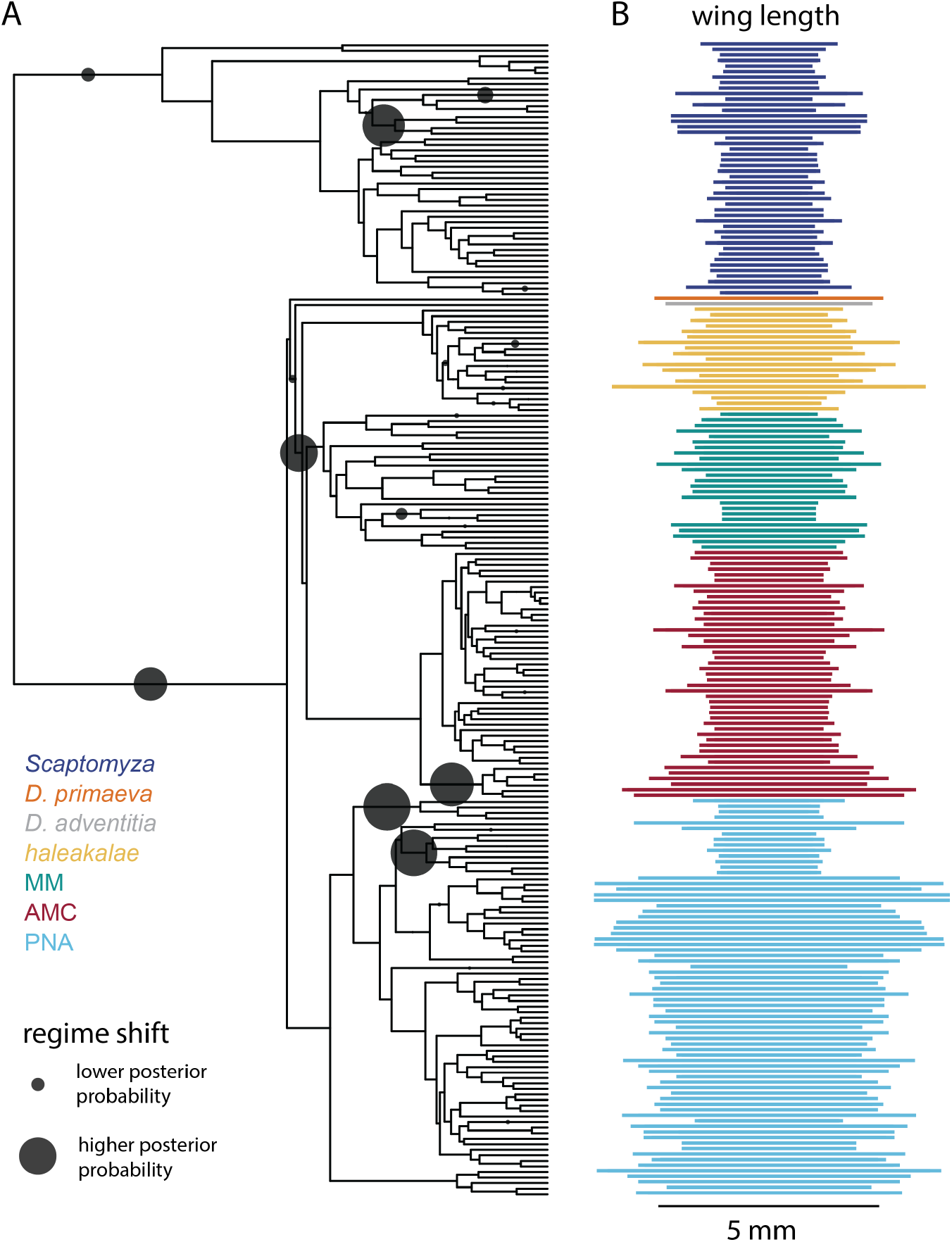
Multiple shifts in evolutionary regimes help explain the diversity of wing length. A, Using the R package bayou^78^, we modeled the evolution of wing length (mm) on the phylogeny and detected several probable shifts in evolutionary regimes (gray circles, larger indicates greater posterior probability that a shift occurred on that branch). Locations of probable shifts include at the base of the AMC+MM+*haleakalae* clade, subtending the *antopocerus* group (AMC), and subtending the *nudidrosophila* (PNA), among others. B, The distribution of wing lengths across the phylogeny of Hawaiian *Drosophila* and *Scaptomyza*.

Over 1000 stochastic character maps, we recovered an average of 44 transitions in oviposition substrate over the evolutionary history of Hawaiian Drosophilidae. The majority of these changes occurred along branches leading to extant tips, with few transitions at internal nodes (on the summary tree, 8 out of 36 total changes). On average, 70% of transitions were between using a single substrate type as a primary host (“specialist” species) and using multiple types (“generalist” species, defined as using any two substrates that each comprise 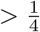 of all rearing records, or with no substrate that comprises 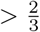 of rearing records^39^.) Other transitions were primarily between using rotting bark, leaves, or sap. Pinpointing branches of likely transitions shows that some groups have experienced many more transitions than others, especially MM and non-flower / spider egg breeding *Scaptomyza*. Most generalist species fall in one of these two clades, which also include specialist bark and leaf breeders, among other substrates.

### Evolution of wing, thorax, and body length

Alongside ecological diversification, the Hawaiian Drosophilidae show substantial diversity in adult body size. We used the time-calibrated phylogeny to model the number and timing of major changes in the evolutionary dynamics of size across the phylogeny. First, we digitized 795 records from 26 publications^24,25,27,37,38,57–77^, including descriptions of body, wing, and thorax length across 552 species. Then we mapped these traits onto our phylogenetic results, and used the R package bayou^78^ to identify branches that represent probable shifts in trait diversification. This package uses Ornstein-Uhlenbeck (OU) models to describe shifts in evolutionary regimes, defined as lineages that share an OU optimum trait value.

In the case of wing length, we find evidence for several highly supported regime shifts in the evolutionary history of Hawaiian Drosophilidae (Fig 5). Some of these are independent shifts on branches subtending groups with larger wings than their nearby relatives, including flies in the *antopocerus* group (AMC) and in the *Engiscaptomyza*+*Grimshawomyia* subgenera (*Scaptomyza*). Others are independent shifts on branches subtending lineages with smaller wings than nearby relatives such as the *nanella*+*ischnotrix* (MM) and the *nudidrosophila* subgroups (PNA). This suggests that the evolutionary history of Hawaiian *Drosophila* has included multiple convergent transitions to both larger and smaller wings. In the case of *nudidrosophila* (PNA), we note that the topology recovered in the summary tree dividing this subgroup into two lineages has very low bootstrap support (Fig. S13, polytomies), and we suggest that the two shifts to smaller wings recovered within PNA may represent a single shift if this group is indeed monophyletic.

We found similar results when considering thorax length (Fig. S15) and body length (Fig: S16). In the case of the former, we find shifts at the base of *antopocerus*, and *nudidrosophila*, consistent with the shifts recovered for wing size. In the case of body size, the most probable shifts are located at the base of the Hawaiian *Drosophila* and the *Engiscaptomyza*+*Grimshawomyia* subgenera. However for body length, no regime shifts received substantially more support than others, despite running bayou for an extra million generations and achieving a final effective size for log-likelihood of 401.9.

### Evidence for convergent evolution of ovariole number and egg size

We also performed these analyses on reproductive traits, including egg size, egg shape (aspect ratio, calculated as egg length / width), and the number of egg producing compartments in the ovary (ovarioles). These traits have been the subject of several life-history studies regarding the hypothesized trade-offs between offspring size and number, and its relationship to ecology^37,37,38,74^. Considering egg shape, we find evidence for a shift at the base of the PNA clade, which have proportionally longer eggs than their relatives (Fig. S17A). In the case of egg volume, we find evidence for independent shifts on branches subtending flies with large eggs (*antopocerus* (AMC) and the *Engiscaptomyza*+*Grimshawomyia*, Fig. S17B). In the case of ovariole number, we find shifts at the base of the *haleakalae*+AMC+MM clades, which have on average fewer ovarioles than the other Hawaiian *Drosophila* (*D. primaeva* and PNA, Fig. S18A).

Work by Kambyesllis and Heed (1971) ^37^ suggested that Hawaiian Drosophilidae species can be grouped into four reproductive categories based on suites of ovarian and egg traits. Subsequent publications^15^, including work by ourselves^38^, showed that these categories largely map to differences in oviposition substrate. Given the evidence that ovary and egg traits may be evolving together, we analyzed them with the R package SURFACE^79^, which uses OU models to analyze regime shifts in multiple traits at once, and allows for distant taxa to share regimes via convergent evolution. The best fitting model indicates four regimes (Fig: S19), two of which correspond to categories defined by Kambysellis and Heed (1971)^37^: [1] very large eggs and low ovariole number in *S. undulata* (*Grimshawomyia*) and *S. nasalis* (*Engiscaptomyza*, group I in their publication); [2] large eggs with moderate to large bodies and moderate ovariole number in *antopocerus* (AMC) and also in *S. crassifemur* (*Engiscaptomyza*) and *S. ampliloba* (*Engiscaptomyza*, group II in their publication). The remaining two regimes redistribute species that fall into groups IIIa and IIIb in Kambysellis and Heed (1971) into groups that have [3] small eggs, moderate to large bodies, and high ovariole number in PNA flies, *D. primaeva* and *D. comatifemora* (MM); [4] flies with small eggs, small to moderate bodies, and moderate ovariole number, in the remaining AMC+MM flies along with *D. preapicula* (PNA) and *S. devexa* (*Elmomyza*). As predicted by Kambysellis and colleagues (1997) ^15^ and ourselves^38^, these final two regimes are largely divided between bark breeding flies (4) and leaf breeding flies (3).

## Discussion

The landscape of treespace, representing support for all the possible topologies given the data, is often hidden from our view^80,81^. This is especially true as the size of datasets grow, making it more laborious to traverse treespace landscapes. Approaches such as visualizing the posterior distribution of parameters in a Bayesian analysis, or alternative hypotheses testing (e.g., an SOWH test in a maximum likelihood framework), can provide a sense of how support for one result compares to others. But given that a complete exploration of treespace is typically not available, we often do not know whether the support landscape in treespace is generally flat, rugged, or highly structured.

Model clades for phylogenetics such as the Hawaiian Drosophilidae, however, offer an opportunity to explore these methods using real-world data. In the case of the landscape of treespace, especially in the context of discordance of gene trees and species trees, these flies have a long history as one such model clade. Here we provide a comprehensive snapshot of treespace for this island radiation. We find that, in this case, the landscape of support is largely defined by one hotspot in both gene and site concordance. This hotspot divides the major clades of Hawaiian *Drosophila* into two main lineages, the picture wing flies and their allies (PNA) on one side, and the *modified-mouthparts* and *modified-tarsus* (AMC) flies on the other. We consider this division to be strongly indicated given the data, and we note that this is in line with other recent phylogenetic results (Fig. S1)^16,17^.

Within this hotspot of support, several alternative topologies that differ in the placement of smaller clades (*D. primaeva* and *haleakalae*) have an equivocal amount of support across genes and sites. We suggest that much of this discordance represents the results of evolutionary processes that took place on the short internodes at the base of the radiation. Despite this local discordance, the outcome of all phylogenetic software tested here indicates strong support for a single topology (Fig. 1A). With this information, we consider that tree, with PNA as the sister clade to the rest of Hawaiian *Drosophila*, and *haleakalae* as the sister clade to AMC+MM, to be a plausible new hypothesis for the evolution of these flies. We suggest that additional taxonomic sampling in the *haleakalae* will be valuable in gaining a fine-scale view of the landscape of support within this hotspot.

This new hypothesis for the relationship between major groups has several implications for our understanding of ecological and morphological evolution. Some previous studies have focused on defining one group as ‘basal’ to others (e.g., *haleakalae*, MM, or *D. primaeva*)^15,16^. However our results provide an alternative interpretation. We find that the PNA clade (including *picture-wing* flies) is the sister clade to all others, and we note that for at least one trait (bark breeding), most PNA flies appear to display the same state as the most common ancestor of Hawaiian *Drosophila*. The relationship between this group to *haleakalae* and others suggests the possibility of a secondary loss of complex courtship behavior in the latter^16^. We note that the overall pattern in the group has been one of many transitions to and from the ancestral state, including in ecology, size, and allometry.

Our results on wing, body, and egg size evolution show that Hawaiian Drosophilidae have experienced multiple, independent shifts to both larger and smaller sizes. These repeated changes present an opportunity to test the predictability of evolution by analyzing whether repeated changes in size are coincident with changes in other features, including ecology, development, and whether these repeated trait changes share the same genetic regulatory basis. The findings of this study on ovariole number and egg size evolution are consistent with what has previously been shown^15,38^, indicating that evolutionary changes in the larval ecology correspond to changes in reproductive trait evolution. However, our findings here show that larval feeding substrate does not explain all the dynamics of trait diversification in Hawaiian *Drosophila*. For example, the *antopocerus* group (AMC) shares the same oviposition substrate as most other AMC flies, yet we find evidence that several important shifts in thorax, wing, and egg size evolution all occurred on the branch subtending its diversification.

Previous authors have commented on the potential of the Hawaiian *Drosophila* as a model clade for the study of the evolution of development^9,35^, given its close relationship to genetic model species like *D. melanogaster*. Progress in this effort has not always been straightforward, however, given their longer generation times and specific host plant requirements to induce oviposition in the lab^9^. We propose that advances in evo-devo study of the Hawaiian Drosophilidae will be added by leveraging evolutionary methods to formulate and test developmental hypotheses. For example, we can use phylogenetic comparative methods to statistically detect signatures of convergent evolution and to identify changes in patterns of allometric growth^34,38^. Going forward, such methods will be essential in providing testable hypotheses regarding the relationship of developmental data to ecological and morphological parameters. The results of these analyses will provide valuable complementary studies to the developmental literature generated using laboratory-amenable model drosophilids, and shed light on the genetic basis of this remarkable island radiation.

## Methods

### Field collection and RNA extraction

#### Field collection

Specimens used for transcriptome sampling were caught on the Hawaiian islands between May of 2016 and May of 2017. Specimens were caught using a combination of net sweeping and fermented banana-mushroom baits in various field sites on the Hawaiian islands of Kaua’i and Hawai’i (see Supplemental Table S1 for locality data). Field collections were performed under permits issued by the following: Hawai’i Department of Land and Natural Resources, Hawai’i Island Forest Reserves, Kaua’i Island Forest Reserves, Koke’e State Park, and Hawai’i Volcanoes National Park. Adult flies were maintained in the field on vials with a sugar-based media and kept at cool temperatures. They were transported alive back to Cambridge, MA where they were maintained on standard *Drosophila* media at 18°C. Samples were processed for RNA extraction between 5 and 31 days after collecting them live in the field (average 9.8 days, see Supplemental Table S1). One species, *Scaptomyza varia*, was caught in the field before the adult stage by sampling rotting *Clermontia sp*. flowers (the oviposition substrate). For this species, male and female adult flies emerged in the lab, and were kept together until sampled for RNA extraction.

#### Species identification

Species were identified using dichotomous keys^19,27,63,70,73^ when possible. Many keys for Hawaiian Drosophilidae are written focusing on adult male specific characters (e.g., sexually dimorphic features or male genitalia)^63^. Therefore, for species where females could not be unambiguously identified, we verified their identity using DNA barcoding. When males were caught from the same location, we identified males to species using dichotomous keys and matched their barcode sequences to females included in our study. When males were not available, we matched barcodes from collected females to sequences previously uploaded to NCBI^16,23,30^.

The following dichotomous keys were used to identify species: for *picture-wing* males and females, Magnacca and Price (2012)^19^; for *antopocerus* males, Hardy (1977)^70^; for *Scaptomyza*, Hackman (1959)^63^; for species in the *mimica* subgroup of MM, O’Grady and colleagues (2003)^73^; for other miscellaneous species, Hardy (1965)^27^.

For DNA barcoding, DNA was extracted from one or two legs from male specimens using the Qiagen DNeasy blood and tissue extraction kit, or from the DNA of females isolated during RNA extraction (see below). We amplified and sequenced the cytochrome oxidase I (COI), II (COII) and 16S rRNA genes using the primers and protocols described in Sarikaya and colleagues (2019)^38^.

For barcode matching, we aligned sequences using MAFFT, version v7.475^82^, and assembled gene trees using RAxML, version 8.2.9^43^. Definitive matches were considered when sequences for females formed a monophyletic clade with reference males or reference sequences from NCBI (Supplemental Table S2). Sequence files and gene trees are available at the GitHub repository, http://github.com/shchurch/hawaiian_drosophilidae_phylogeny_2021, under `analysis/data/DNA_barcoding`.

Female *D. primaeva, D. macrothrix, D. sproati*, and *D. picticornis* could be identified unambiguously using dichotomous keys. Female *D. atroscutellata, D. nanella, D. mimica, D. tanythrix, S. cyrtandrae, S. varipicta*, and *S. varia* were identified by matching barcodes to reference sequences from NCBI, reference males, or both. For the female *haleakalae* fly used in this study, no male flies were caught in the same location as these individuals, and no other sequences for *haleakalae* males on NCBI were an exact match with this species. Given its similar appearance to *Drosophila dives*, we are referring to it here as *Drosophila* cf *dives*, and we await further molecular and taxonomic studies of this group that will resolve its identity. Photos of individual flies used for transcriptome sequencing are shown in Fig. S20.

#### RNA extraction

RNA was extracted from frozen samples using the standard TRIzol protocol (http://tools.thermofisher.com/content/sfs/manuals/trizol_reagent.pdf). One mL of TRIzol was added to each frozen sample, which was then homogenized using a sterile motorized mortar. The recommended protocol was followed without modifications, using 10 µg of glycogen, and resuspending in 20µL RNAse-free water-EDTA-SDS solution. DNA for subsequent barcoding was also extracted using the phenol-chloroform phase saved from the RNA extraction.

RNA concentration was checked using a Qubit fluorometer, and integrity was assessed with an Agilent TapeStation 4200. RNA libraries were prepared following the PrepX polyA mRNA Isolation kit and the PrepX RNA-Seq for Illumina Library kit, using the 48 sample protocol on an Apollo 324 liquid handling robot in the Harvard University Bauer Core Facilities. Final library concentration and integrity were again assessed using the Qubit and TapeStation protocols.

The field collecting for this study was accomplished with a target number of individuals per species in mind, based on future sampling objectives for RNA sequencing studies that, as of the time of writing, have not been published. These objectives were to have four wild-caught, mature, apparently healthy females, three of which were to be dissected for tissue-specific RNA sequencing, and one intended as a whole-body reference library. When four individuals were not available, the reference library was assembled by combining the tissue specific libraries from one of the other individuals. This was the case for the following species: *D. sproati*, which was dissected and had RNA extracted separately from the head, ovaries, and carcass, with RNA combined prior to library preparation; and *S. varia, S. cyrtandrae* and *D*. cf *dives*, for which RNA was extracted and libraries prepared for separate tissues, and raw reads were combined after sequencing.

For the other eight species, sufficient individual females were available such that reads for transcriptome assembly were sequenced from a separate individual. In these cases one entire female fly was dissected and photographed to assess whether vitellogenic eggs were present in the ovary, and all tissues were combined in the same tube and used for RNA extraction.

Libraries for transcriptome assembly were sequenced on an Illumina HiSeq 2500, using the standard version 4 protocol, at 125 base pairs of paired-end reads. A table of total read counts for each library can be found in Supplemental Table S3.

### Transcriptome assembly

Transcriptome assembly was performed using the agalma pipeline, version 2.0.0^46^. For the 12 new transcriptomes presented in this study, reads from separate rounds of sequencing were concatenated and inserted into the agalma catalog. These were combined with seven publicly available outgroup genomes (*D. virilis, D. mojavensis, D. pseudoobscura, D. ananassae, D. willistoni*, and *D. melanogaster* ^41^), two Hawaiian genomes (*D. grimshawi*^41^ and *D. murphyi*^40^), and four *Scaptomyza* genomes (*S. graminum, S. montana, S. hsui, S pallida*^40^). For the non-Hawaiian *Drosophila* and *D. grimshawi* genomes, the longest isoform per gene was selected using the gene header. For the four *Scaptomyza* genomes and *D. murphyi* genomes, single copy orthologs were filtered using BUSCO version 4.1.4^83^ against the Diptera obd10 gene set (over 98% of genes were retained as single-copy orthologs). See Supplemental Table S4 for genome information.

Using the agalma pipeline, the quality score of each library was assessed, and transcriptomes were assembled using the standard parameters. The publicly available genomes were translated and annotated, and the homology of assembled products was inferred using the all-by-all blast component of the homologize step in the agalma pipeline, using nucleotide data and a GTR+Gamma model to infer gene trees. The agalma version 2.0.0 pipeline also performs a step to reduce the effects of transcript misassignment using a phylogenetically informed approach (treeinform)^84^. Gene orthology was inferred according to the topology of gene trees estimated with RAxML, orthologs were aligned and trimmed using MAFFT^82^ and Gblocks^85^ respectively, and a supermatrix of aligned orthologous sequences was exported.

The final supermatrix output from agalma consisted of 10,949 putatively orthologous genes and 12,758,237 sites. For the primary analyses performed in this manuscript, this supermatrix was not filtered by occupancy, and the actual gene occupancy was 41.9% across the 24 species present in this study. We also created a supermatrix filtered to a target occupancy of 80%, which consisted of 1,926 genes and 1,943,000 sites, which we used to reestimate the maximum likelihood phylogeny.

All commands used to run the agalma pipeline, and all output report files, are available at the GitHub repository http://github.com/shchurch/hawaiian_drosophilidae_phylogeny_2021, `under analysis/phylotranscriptomics`.

### Phylotranscriptomics and concordance factors

We estimated the maximum likelihood phylogeny using IQtree, version 2.1.1^42^. We ran IQtree on a dataset partitioned by transcripts, and using the default Model Finder^86^ per partition^87^. For this analysis, partitions containing no informative sites were excluded. We estimated 1000 ultrafast bootstraps^88^ for each node. We also used IQtree to estimate the gene and site concordance factors, as described in Minh and colleagues (2020)^47^. We ran this analysis first on a concatenated dataset output by the agalma pipeline command supermatrix with no gene occupancy threshold (returning all aligned transcripts), and then repeated it on a matrix with an 80% occupancy threshold. All subsequent phylogenetic analyses were performed on the full dataset.

We also estimated the maximum likelihood phylogeny using the speciestree step of the agalma pipeline, which itself runs RAxML, version 8.2.9^43^. We used the default parameters for RAxML as called within the agalma phylotranscriptomic pipeline (model GTR+Gamma, 1000 bootstraps).

We compared the most likely tree against two alternative hypotheses (Fig. 1B-C) using the Swofford-OlsenWaddell-Hillis (SOWH) test^2^, as implemented in sowhat, version 1.0^89^. We ran both comparisons using a GTR+Gamma model, unpartitioned data file, and 100 simulated datasets.

We estimated the phylogeny using the Bayesian software PhyloBayes, mpi version 1.7a^44^. We ran PhyloBayes using a CAT-GTR model for nucleotide data, without partitions, on the full set of transcripts exported from agalma. Phylobayes was run for 1400-1900 generations, and convergence was assessed as the maximum difference between two chains. The initial two chains did not show signs of convergence after 1000 generations (maximum difference was 1), so two additional chains were initialized. These reached convergence with one of the initial chains at 450 generations (maximum difference was 0). The divergence between these three chains and the fourth resulted from differences in the relationships between the MM, AMC, and *haleakalae* clades. The consensus tree was estimated using all four chains and burn-in of 100 generations, taking every tree (maximum difference was 1).

We estimated the phylogeny using the multi-species coalescent with the software ASTRAL, version 5.7.7^45^. For this analysis we input the gene trees as inferred by IQtree, using the methods described above.

To further explore the concordance of data across possible topologies in treespace, we wrote a custom python script to create all 105 combinations of possible topologies for the five clades in question, with the root between these clades set at the split between Hawaiian *Drosophila* and *Scaptomyza*. We used each of these trees as the constraint for a concordance factor analysis, using the same approach as described above for the most likely tree. We visualized treespace by plotting each tree according to Robinsoun-Foulds distance using the R package treespace, version 1.1.4^90^. We then mapped on this space the mean concordance factors for each topology (calculated as the mean site and gene concordance on branches, excluding those shared between all topologies).

We performed likelihood mapping analyses on each partition from the full supermatrix, using the lmap command in IQtree2 and using all possible quartets. For each partition, we performed two analyses, first analyzing the arrangement of the PNA clade relative to *Scaptomyza, D. primaeva*, and a clade uniting AMC+MM+*haleakalea*, and second, analyzing the arrangement of the *haleakalae* clade relative to MM, AMC, and a clade uniting *PNA* and *D. primaeva*. We summarized all likelihood mapping results by counting the total number of quartets across all partitions that fell into each area of a three- and seven-way partitioned likelihood map^48^.

We calculated a conservation score for each gene in our supermatrix, using the software trimAL, version v1.4.rev22, as the accumulated similarity score divided by the number of residues in each alignment, following the protocol of Cunha and colleagues (2021)^91^.

We tested the robustness of results to potential errors in multiple sequence aligment by filtering out poorly aligned sequence fragments and repeating the inference and concordance analyses. Poorly alignment fragments were detected and removed using the software spruceup, version 2020.2.19^92^, with default parameters and cutoffs at 0.95, 0.97, and 0.99 with which outlier fragments were removed.

All commands used to execute the concordance factor analysis are included in the GitHub repository http://github.com/shchurch/hawaiian_drosophilidae_phylogeny_2021 in the rmarkdown file for the supplement of this manuscript, as well as the folder `analysis/phylotranscriptomics/concordance-factor`.

### Estimating an expanded phylogeny

We used the phylotranscriptomic results above, combined with previously published genetic data for Hawaiian Drosophilidae, to estimate an expanded phylogeny. First we gathered the accession numbers from all previously published studies of Hawaiian Drosophila and Scaptomyza genetics^10,17,23,29,29–33^. Nucleotide data for each accession number were downloaded from NCBI in March of 2019. We made no manual alterations to these sequences, with the following exceptions: We replaced all non-nucleotide sites (e.g., ‘N’, ‘R’) with missing data (‘?’); we removed two sequences (U94256.1 - *D. disjuncta* and U94262.1 - *S. albovittata*) from the 16S dataset that did not align to other sequences; we manually removed a portion of the COI dataset that did not align; we corrected spelling for *S. albovittata*; and we updated the taxonomic name of *D. crassifemur* to *S. crassifemur*. Original and modified sequences are provided at the GitHub repository http://github.com/shchurch/hawaiian_drosophilidae_phylogeny_2021 under `analysis/time-calibrated_phylogenetics/downloaded_sequences`.

We then aligned these sequences using MAFFT, version 7.457^82^, --auto option. We visualized alignments, and for gene 16S we repeated the alignment using the --adjustdirectionaccurately option. We removed all information from the headers with the exception of the species name, and then selected the sequence per species with the fewest gaps. We concatenated sequences using phyutility version 2.2.6^93^.

Using these concatenated sequences, we estimated a phylogeny for 316 species, including 271 described species and 45 that are undescribed but for which genetic vouchers were available. This tree was estimated using IQtree^42^ with the topology constrained using the most likely phylotranscriptomic tree. This constraint tree included only taxa overlapping between the phylotranscriptomic and concatenated datasets, with one exception: *D. iki* was substituted for *D*. cf *dives*, given that this unidentified species was the only representative from the *haleakalae* clade present in the phylotranscriptomic analysis. No partition model was used for this analysis. We ran IQtree with default parameters, and we estimated 1000 ultrafast bootstrap support values as well as 1000 SH-like likelihood ratio tests.

Visualizing the results showed that all major clades (AMC, PNA, MM, *haleakalae*, and *Scaptomyza*) were recovered as monophyletic, with the exception of the placement of *D. konaensis*, a member of the hirtitibia subgroup that was recovered as the sister taxon to the AMC clade. We investigated the source of this discrepancy by analyzing the individual gene trees that had representation for this species (COI, COII, and 16S, tree estimated using IQtree as described above, tree files available at `analysis/time-calibrated_phylogenetics/iqtree/iqtree_investigations`). These gene trees showed that *D. konaensis* sequences had variable affinity to unlikely relatives, including *Scaptomyza* and *modified-mouthpart*. We considered this to be an artifact of a possible error in accession sequence, and so we removed *D. konaensis* from downloaded sequences and repeated the alignment and tree estimation steps.

All commands used to download and align sequences as well as estimate the phylogeny, along with all input and output files, are available at the GitHub repository http://github.com/shchurch/hawaiian_drosophilidae_phylogeny_2021, under `analysis/time-calibrated_phylogenetics/`.

### Calibrating the phylogeny to time

This expanded phylogeny was calibrated to time using BEAST, version 2.6.3^49^. This tree search was performed using the following parameters and priors, set using BEAUTi2^94^: a relaxed log-normal clock model, a general time reversible (GTR) site model with 4 gamma categories, and a Yule process branching model. We tested three calibration schemes based on previously published analyses: [1] Four normally distributed node priors, used by Magnacca and Price (2015)^17^, based on the apparent progression rule seen in the island distribution of these species; [2] the same four node priors, but with island ages adjusted to correspond to recently updated estimates for the age at which islands became habitable ^51^, which are based on models that describe the volcanic growth and decay of each Hawaiian island as it has passed over the tectonic hotspot; and [3] four node priors based on progression rule island ages and one node prior based on a single fossil specimen in dominican amber (*S. dominicana*), used by Russo and colleagues (2013)^52^. Calibration times are shown in Table 1.

For all BEAST analysis, the most likely topology from the expanded IQtree search was used to create a starting tree, rooted at the split between *Scaptomyza* and *Drosophila* and with branch lengths removed. This topology was fixed throughout the analysis by setting tree topology operator weights to zero.

The BEAST analyses using only island-age based calibrations were run between 20-25 million generations. The maximum clade credibility tree was determined using TreeAnnotator^95^, with a burn-in of 10%, selected by visualizing in Tracer, version 1.7.1^96^. The analysis run using both island-age and the single fossil node calibration was run 50 million generations, and summarized with a burn-in of 5%. For all analyses, the effective size for the posterior was >100 (Magnacca calibration = 581.3, Magnacca updated island ages = 453.7, Russo calibration island ages = 168.7), though for tree height the effective size for the older island ages and fossil calibrated analyses were both below 100 (Magnacca calibration = 137.4, Magnacca calibration, updated island ages = 92.1, Russo calibration = 87.7).

### Estimating ecological and morphological evolutionary transitions

For ecological data on oviposition and larval feeding substrate, we used the rearing records summarized in Magnacca and colleagues (2008)^39^, Appendix I. Following the method of Magnacca *et al*., we grouped oviposition substrates into several general categories, listed in Supplemental Table S5. We also followed the definition from Magnacca and colleagues of non-monophagous (here referred to as generalist) as any species for which no single substrate type comprised more than 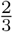 of rearing records, or for which any two substrates each comprised more than 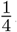. We note that *D. comatifemora* was listed as a bark breeder in Sarikaya and colleagues (2019)^38^, but no rearing records for this species are present in Magnacca and colleagues 2008^39^and Magnacca and O’Grady (2009) ^76^ list it as “breeding habits unknown”.

We reconstructed the ancestral state for general oviposition substrate type using the R package phytools, version 0.7-70^97^ on the maximum clade credibility tree from the constrained BEAST analyses. We performed 1000 simulations of stochastic character mapping using the `make.simmap` function (with a maximum likelihood method for estimating the transition model), and then summarized the ancestral state at each node as the oviposition substrate with the highest posterior probability. We used this summary tree to identify branches with likely transitions between oviposition substrates.

For morphological data on wing, body, and thorax length, we digitized data from 26 publications^24,25,27,37,38,57–77^. For data on ovariole number, egg width, and egg length, we digitized data from three publications^37,38,74^. We made the following modifications to morphological data: In the data from the GitHub repository associated with the study by Sarikaya and colleagues (2019)^38^, egg size was measured using the radius rather than the diameter; therefore for consistency across studies, we multiplied the reported egg measurements by two. We also excluded data on the egg size of *D. incognita* from the same publication^38^ which had two measurements that showed significantly more variation than other measurements (∼150% discrepancy in egg length). We excluded data on wing and thorax length from the manuscript by O’Grady and colleagues (2003) revising the *mimica* subgroup^73^, for which all data were inconsistent with measurements from conspecific species in other studies, possibly due to differences in the measurement method.

We identified shifts in evolutionary regimes using the R packages bayou, version 2.2.0^78^ and SURFACE, version 0.5^79^ on the maximum clade credibility tree from the constrained BEAST analyses. For all analyses, trait data were log_10_ transformed. For species that had multiple records for the same trait across publications, we randomly selected one description (data on intraspecific variation or measurement error were not included in analyses due to inconsistency in the methods used to gather and report these data by the original authors). The bayou analyses were performed using a half-Cauchy distribution for the prior value of alpha and sigma^2^ (scale set at 0.1), a conditional Poisson distribution for the number of shifts (lambda of 10, max of 50), and a normal distribution for theta values (prior mean and standard deviation set at the mean and standard deviation of the trait data). These analyses were run for one million generations, with the exception of body and thorax length, which were run for two million generations. A burn-in value was set at 0.3 and convergence was evaluated using effective size of the likelihood and the number of shifts (k), see Supplemental Table S6. The SURFACE analyses were performed on a combination of egg volume, ovariole number, and body length using default parameters and using a two-step forward-backward process of selecting the number of regimes^79^.

## Data availability

All data, code, tree files, and other results are available at the GitHub repository http://github.com/shchurch/hawaiian_drosophilidae_phylogeny_2021, commit b12cbb10. This code was implemented in a clean computational environment, which can be recapitulated by following the document `build_conda_environment.sh`. Code to reproduce the figures and text for these mansucript files are available as rmarkdown documents. Concordance value results for each of the possible topological arrangements are available at the above GitHub repository. Raw RNA sequencing data are available at the Sequence Read Archive of NCBI, under BioProject PRJNA731506. Assembled transcriptomes and DNA barcode sequences are available at the above GitHub repository.

## Acknowledgments

This work was partially supported by National Science Foundation Graduate Research Fellowship Program DGE1745303 to SHC, National Institutes of Health award R01 HD073499-01 (NICHD) to CGE, and funds from Harvard University to support SHC and CGE. We thank Didem Sarikaya, Karl Magnacca, and Steve Montgomery for their field assistance and expertise in Hawaiian fly identification and husbandry. We thank Kenneth Kaneshiro for the use of his lab space in preparing wild caught specimens. We thank Tauana Cunha and Bruno de Medeiros for providing scripts to run phylogenetic software on Harvard’s computing cluster, and Casey Dunn for providing scripts to handle large genetic data files. We thank members of the Extavour Lab for discussion of ideas, and anonymous reviewers for their helpful suggestions.

