## Supplementary Methods for "Phylotranscriptomics reveals discordance in the phylogeny of Hawaiian *Drosophila* and *Scaptomyza* (Diptera: Drosophilidae)"

### Previous phylogenetic hypotheses

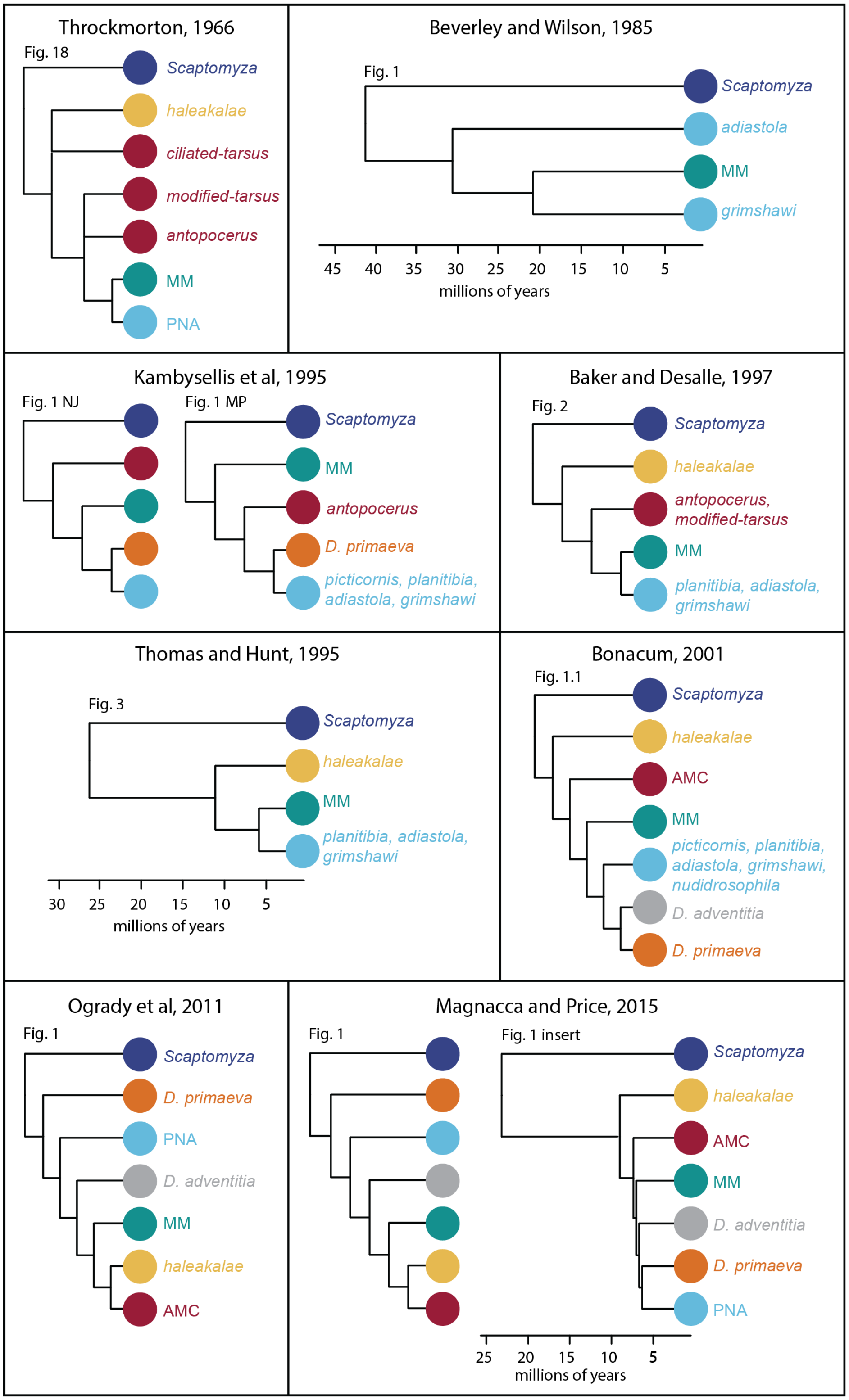

Figure S1: Selected previously published phylogenetic hypotheses for the relationships between clades of Hawaiian Drosophilidae. Figure labels indicate the figure number as originally published[1–8].

### Phylotranscriptomic results

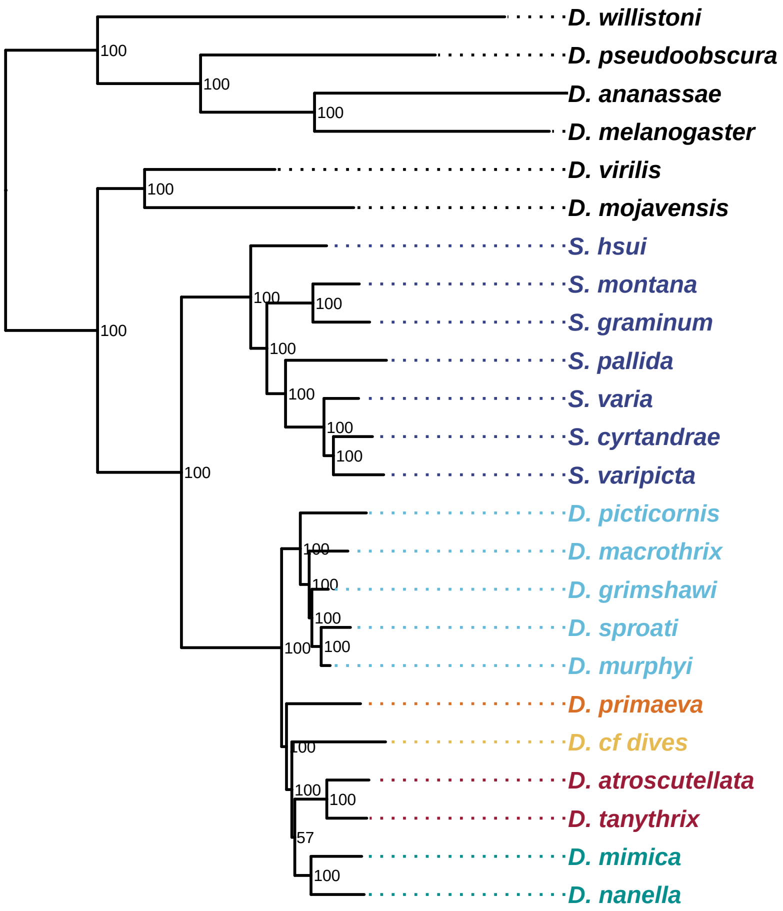

Figure S2: Most likely tree estimated using RAxML. Node labels show bootstrap values. Colors correspond to clades described in Fig. 1 and S1.

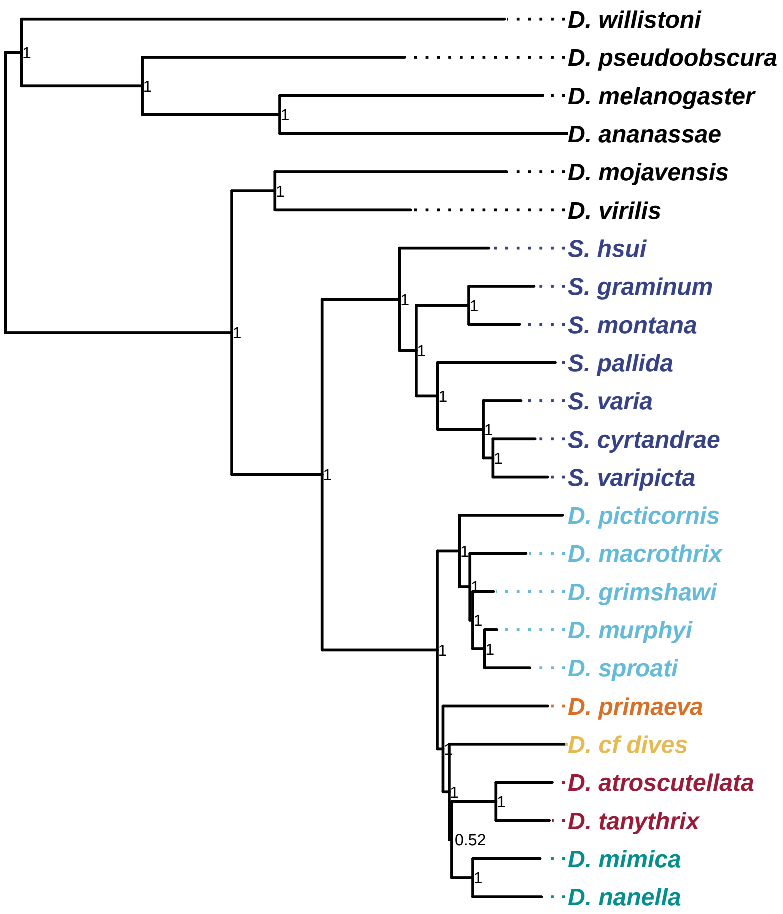

Figure S3: Consensus tree estimated using PhyloBayes. Node labels show posterior support. Colors correspond to clades described in Fig. 1 and S1.

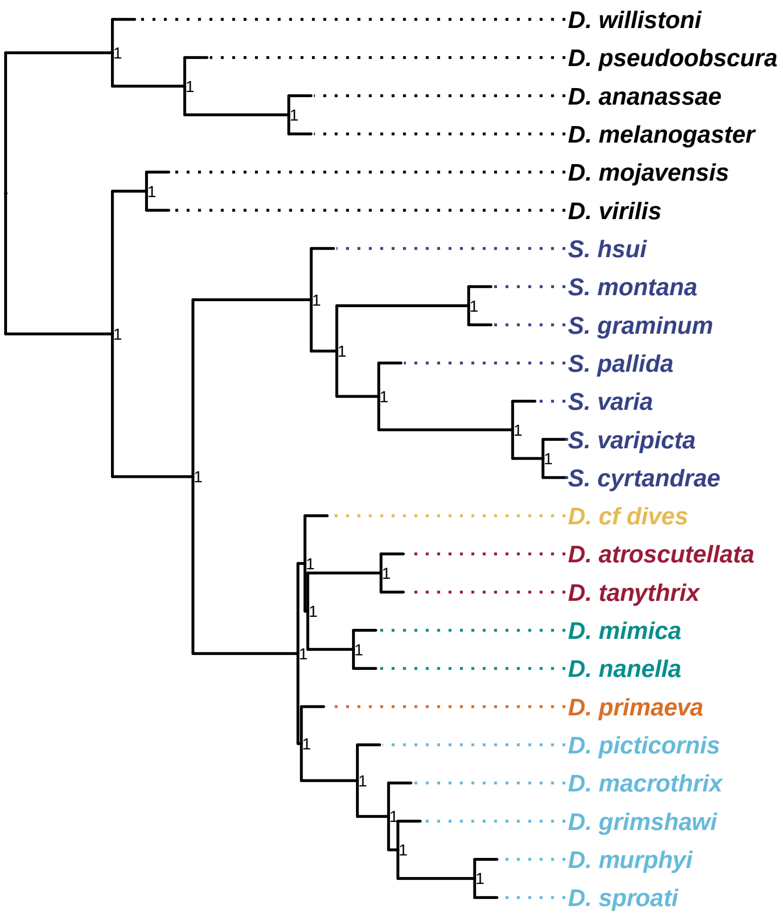

Figure S4: Coalescent tree estimated using ASTRAL. Node labels show local posterior probabilities. ASTRAL estimates branch lengths for internal nodes only, therefore tip branch lengths have been artificially set to a length of 0.5 coalescent units. Colors correspond to clades described in Fig. 1 and S1.

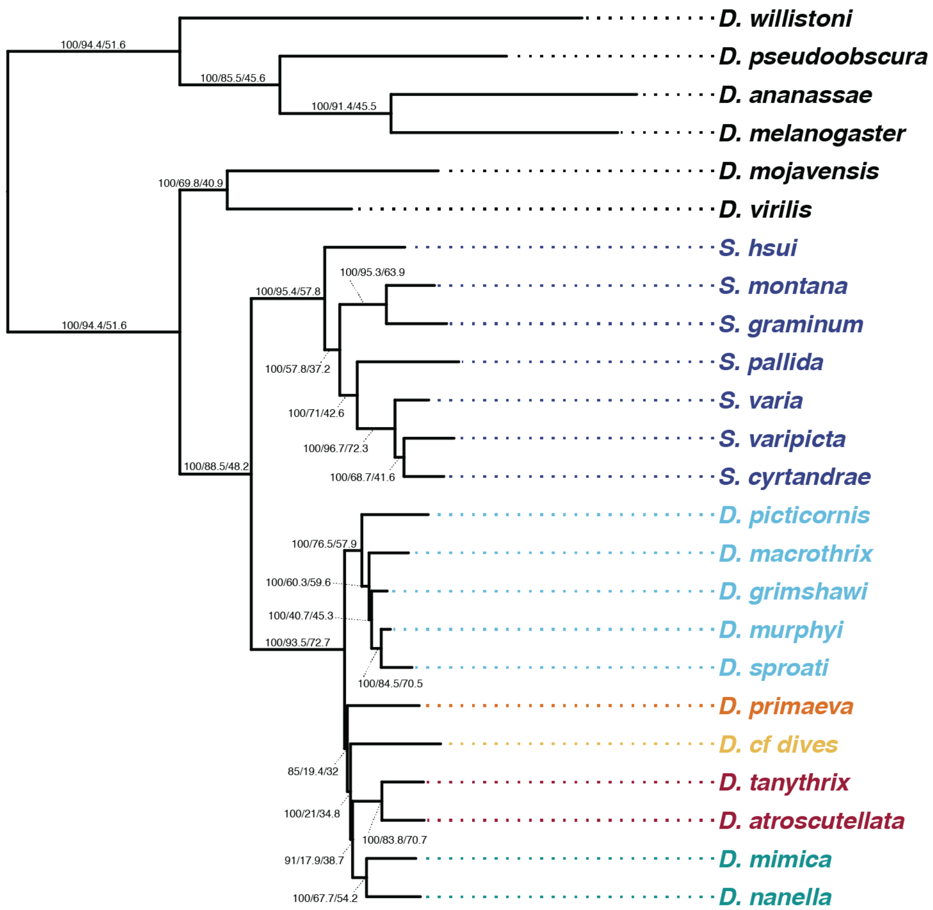

Figure S5: Most likely topology estimated using IQtree on a trimmed dataset, using a an occupancy threshold of 80%. Node labels show boostrap values / gene concordance factors / site concordance factors. Colors correspond to clades described in Fig. 1 and S1.

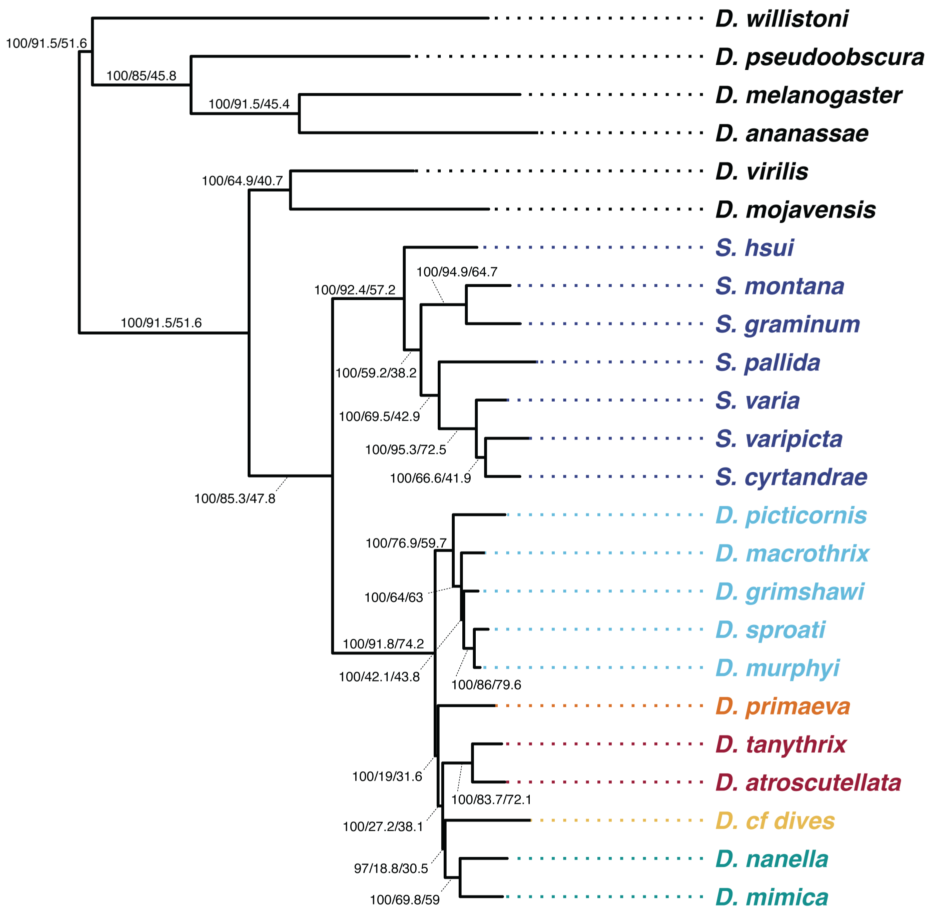

Figure S6: Most likely topology estimated using IQtree on a dataset filtered to exclude poorly aligned sequence fragments, above a 95% cutoff. Node labels show boostrap values / gene concordance factors / site concordance factors. Colors correspond to clades described in Fig. 1 and S1.

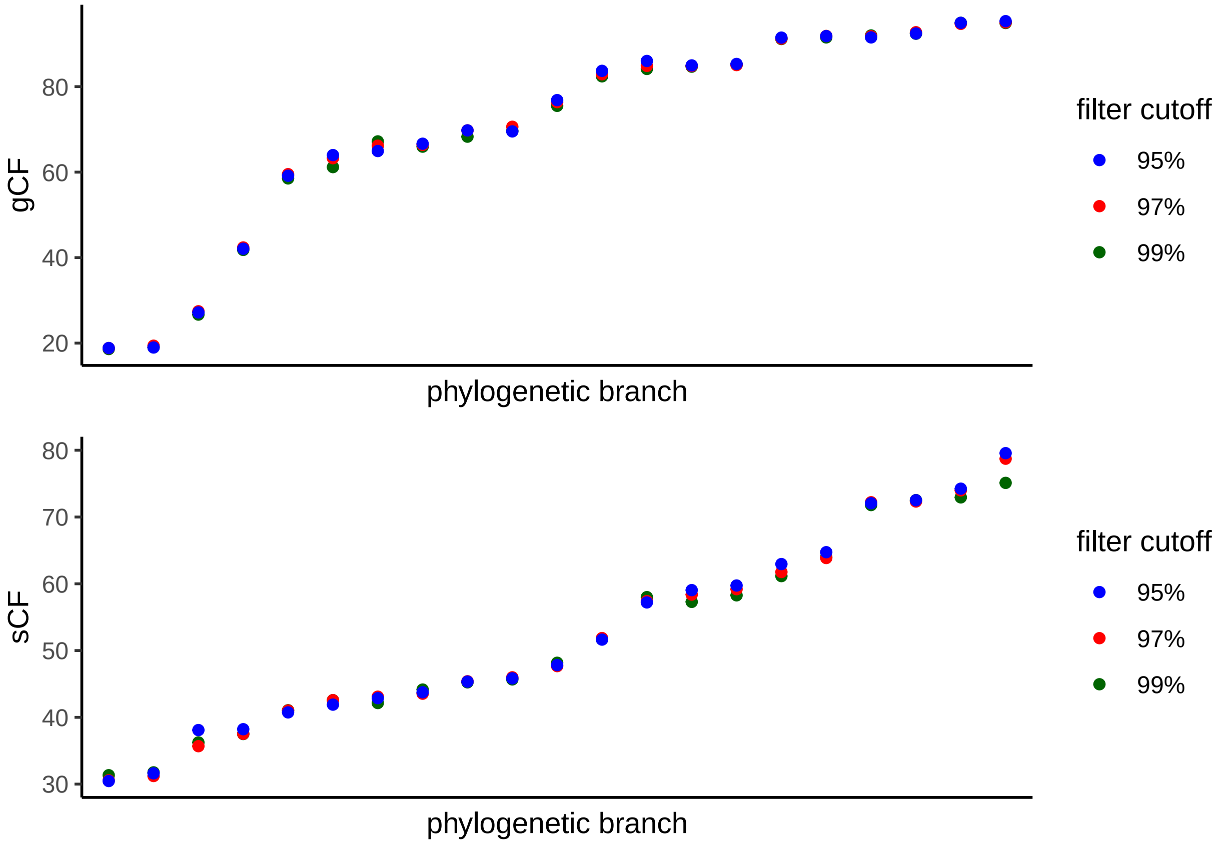

Figure S7: Comparison of concordance factors across branches using variable cutoff values to filter potentially poorly aligned sequence fragments. In both panels, the x-axis shows the 21 branches of the phylogeny, arranged from lowest concordance factor value to highest.

### Concordance factor analysis

There are 210 unique internal branches across all possible topologies of Hawaiian *Drosophila* clades, when the root of the phylogeny is considered to be fixed at the base of the split between *Scaptomyza* and Hawaiian *Drosophila*, and each of the major clades is considered to be monophyletic. Each of these branches defines a relationship between four groups , and in rooted trees like those considered here, one of those groups includes the outgroup. These 210 branches can be divided into four categories:

[1] 15 branches that define the split between *Scaptomyza* and Hawaiian *Drosophila*, which differ based on the arrangement of clades on the *Drosophila* side of the branch. These have universally high gene tree concordance (minimmum of 89.38), and the small amount of variation between them can be attributed to variation in the number of informative sites.

[2] 70 branches that define a relationship that unites any two clades on one side of a branch (panel A in Figs. S8 and S9). These branches indicate support for two clades as sister to one another, and variation across these branches shows that more genes support the unification of *D. primaeva*+PNA and any two of the clades AMC, MM, and *haleakalae*, relative to other groupings.

[3] 90 branches that define a split between two clades of Hawaiian *Drosophila* and the other three (Figs. S8 and S9, panel B). Variation in support across these branches shows a marked increase in the number of genes that support AMC+MM+*haleakalae*, relative to other groupings.

[4] 35 branches that define the split at the base of the Hawaiian *Drosophila* as having one clade sister to the rest of Hawaiian *Drosophila* (or in other words, a branch separating one clade from the other four, Figs. S8 and S9, panel C). Variation in support across these branches shows that the fewest number of genes support either MM or AMC as sister to the rest.

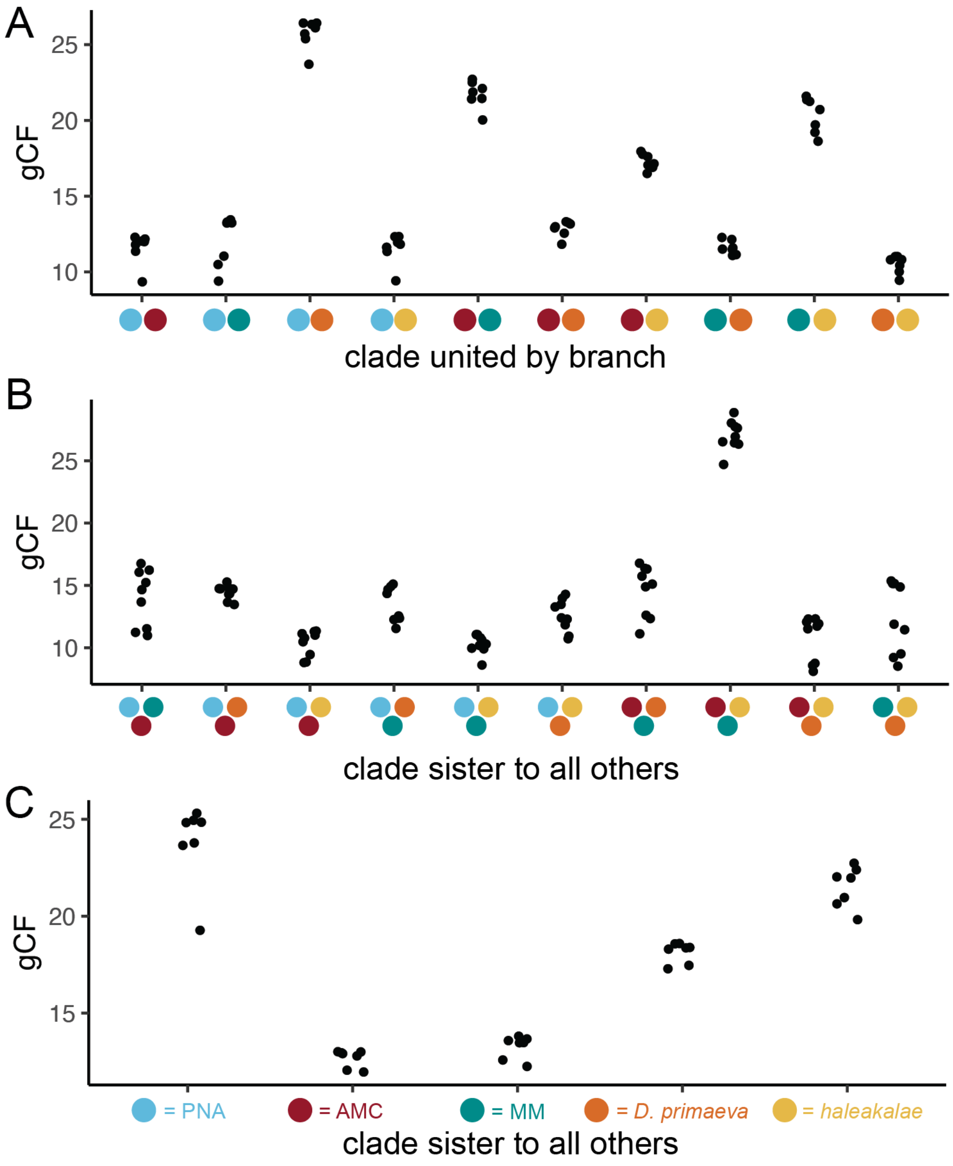

Figure S8: Gene concordance (gCF) across all possible branches.

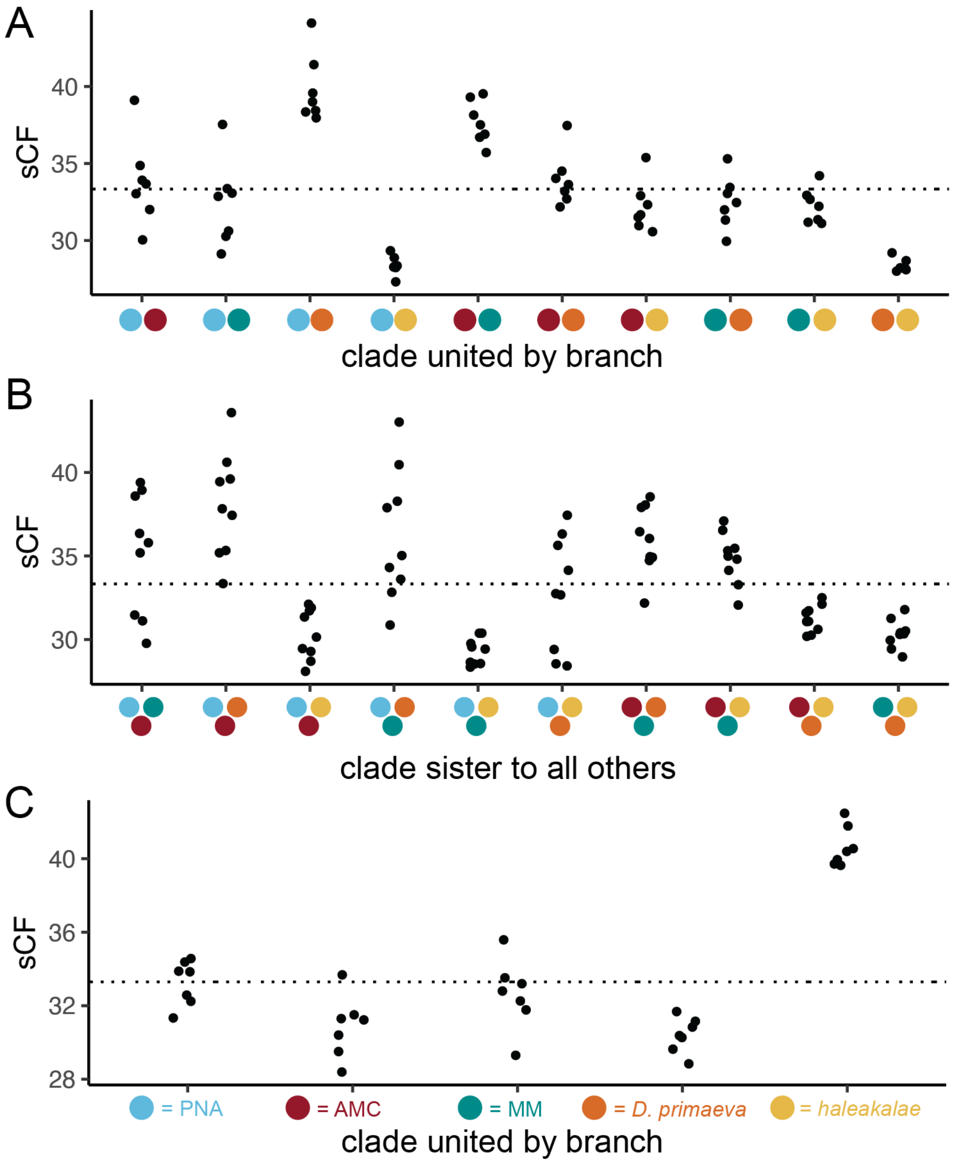

Figure S9: Site concordance (sCF) across all possible branches.

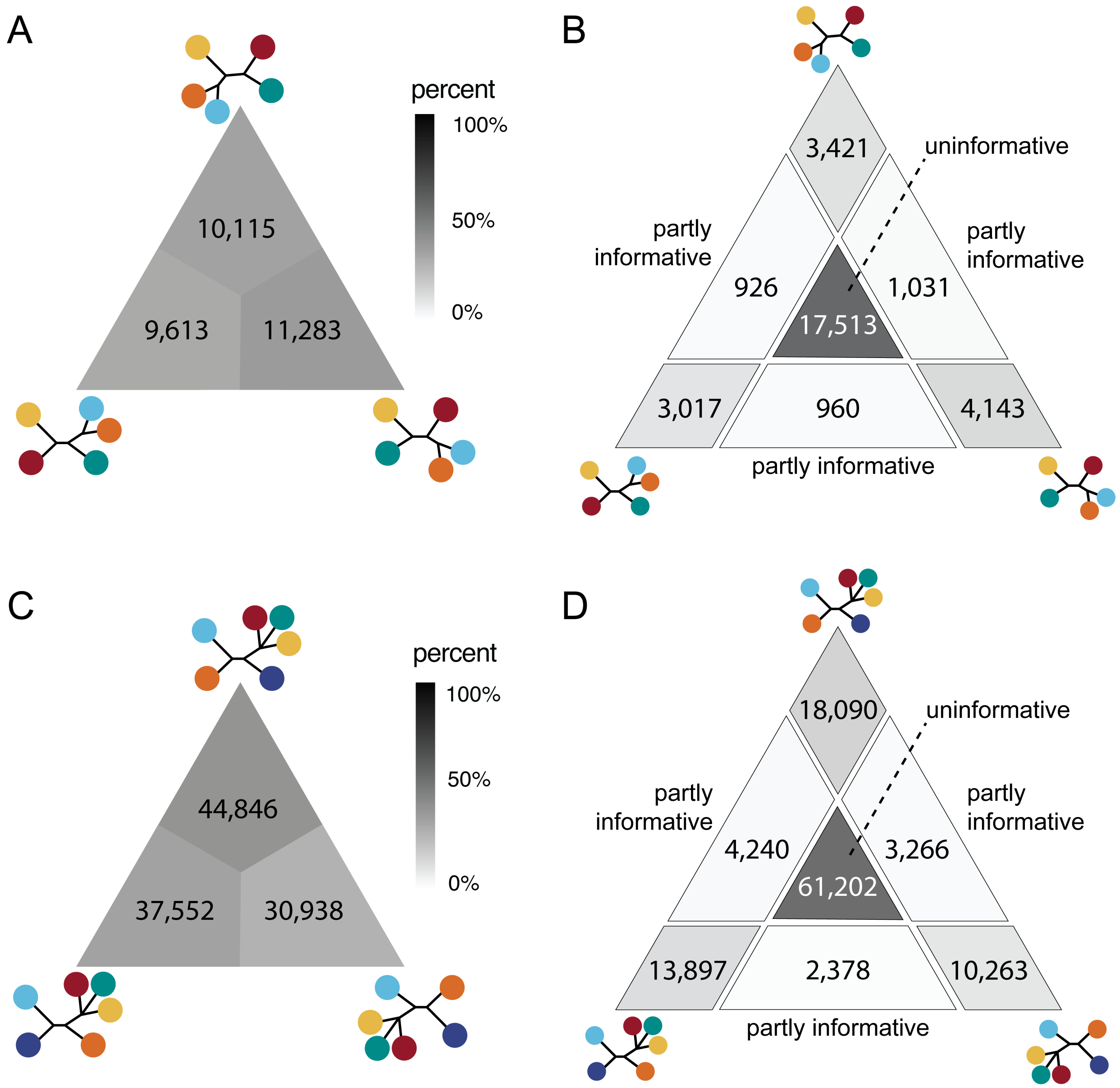

Figure S10: Results of a likelihood mapping analysis of all taxon quartets across all genes. A-B show support for the branches subtending the *haleakalae* group, relative to AMC, MM, and PNA+*D. primaeva*. C-D show support for the branches subtending the PNA group, relative to *D. primaeva*, AMC+MM+*haleakalae*, and *Scaptomyza*. Panels A and C show the number of taxon quartets that support the arrangements shown at the vertices. Panels B and D show the number of quartets that are clearly informative for these arrangements, compared to quartets that are only partly informative or uninformative in distinguishing these arrangements. Triangle panels are shaded by the percent of quartets.

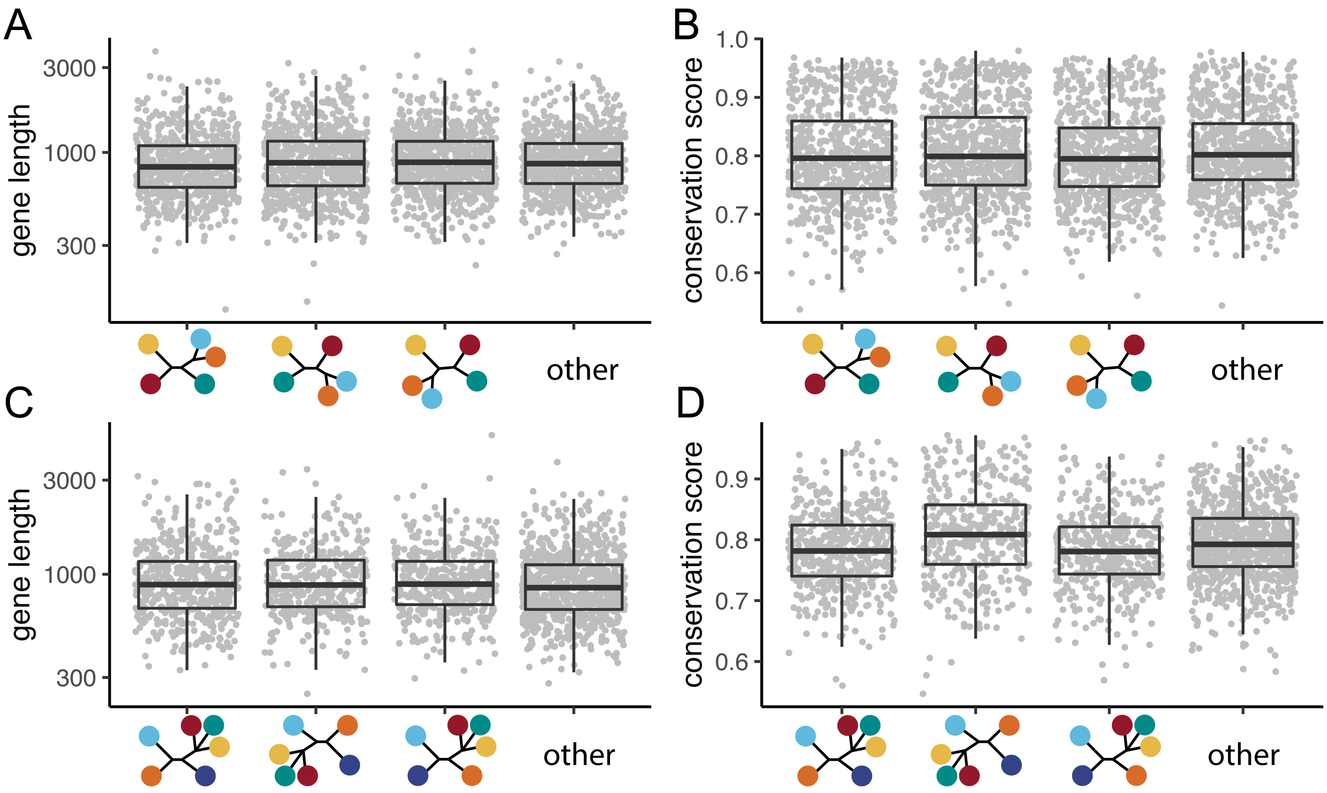

Figure S11: Distributions of genes supporting particular arrangements by gene length and evolutionary conservation. A-B show support for the branches subtending the *haleakalae* group, relative to AMC, MM, and PNA+*D. primaeva*. C-D show support for the branches subtending the PNA group, relative to *D. primaeva*, AMC+MM+*haleakalae*, and *Scaptomyza*. Boxplots show the median, quartiles, and outlier values.

### Expanded phylogenetic analysis

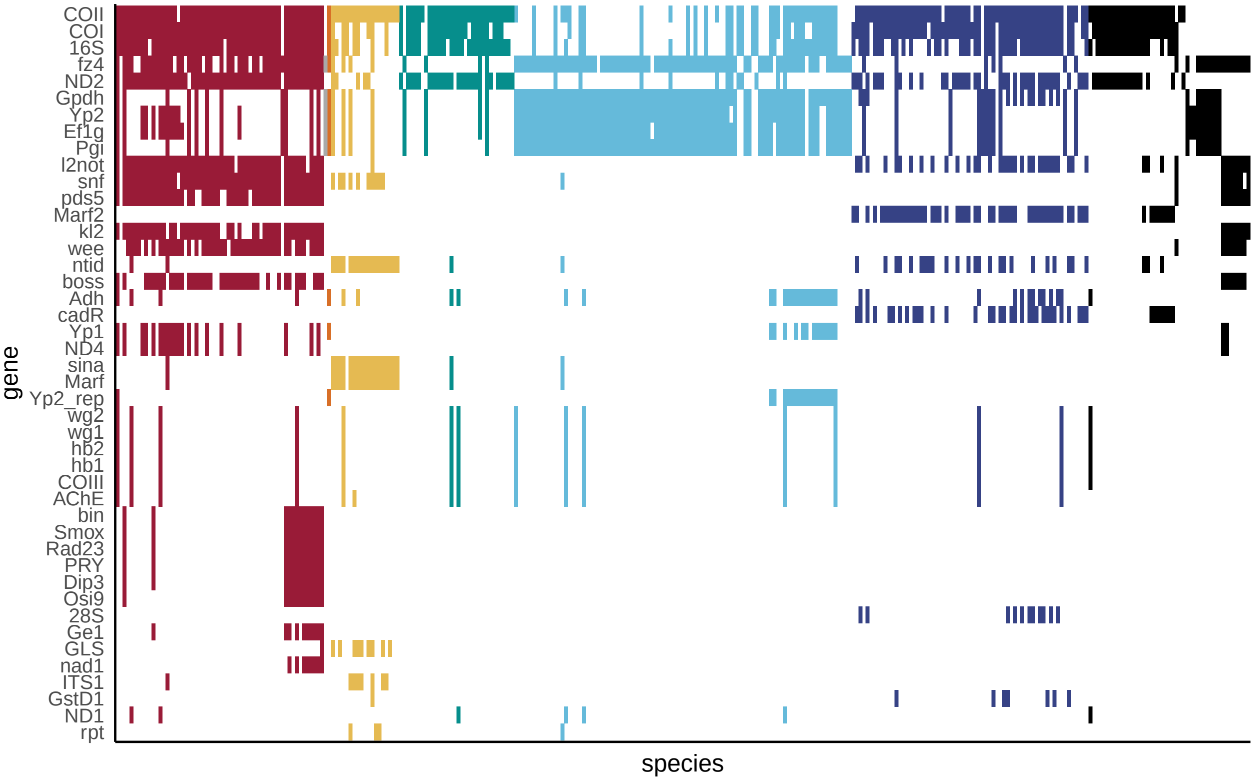

Figure S12: Occupancy matrix of genes in the expanded phylogenetic analysis using previously published mitochondrial and nuclear genetic data, ordered by high to low occupancy on the y axis and by clade, subgenus, group, and subgroup on the x axis. Colors correspond to Fig. 1 and S1, black indicates undescribed species.

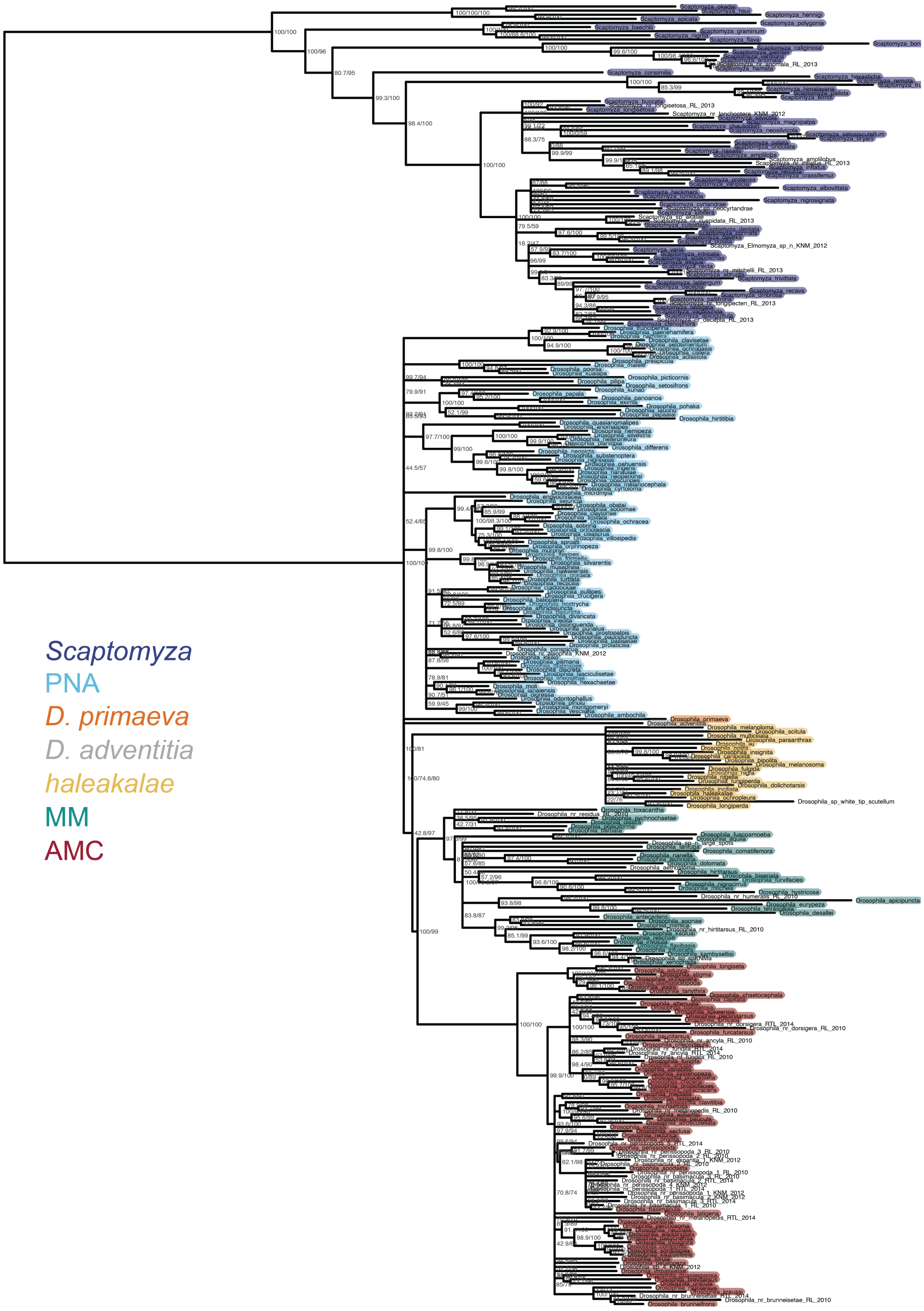

Figure S13: Most likely tree estimated with IQtree using previously published genetic data. The tree search was constrained to follow the relationships estimated using phylotranscriptomic data. Support values shown are SH-like approximate likelihood ratio test / ultrafast bootstrap. Nodes with an ultrafast bootstrap support <95 have been collapsed.

### Oviposition ecology

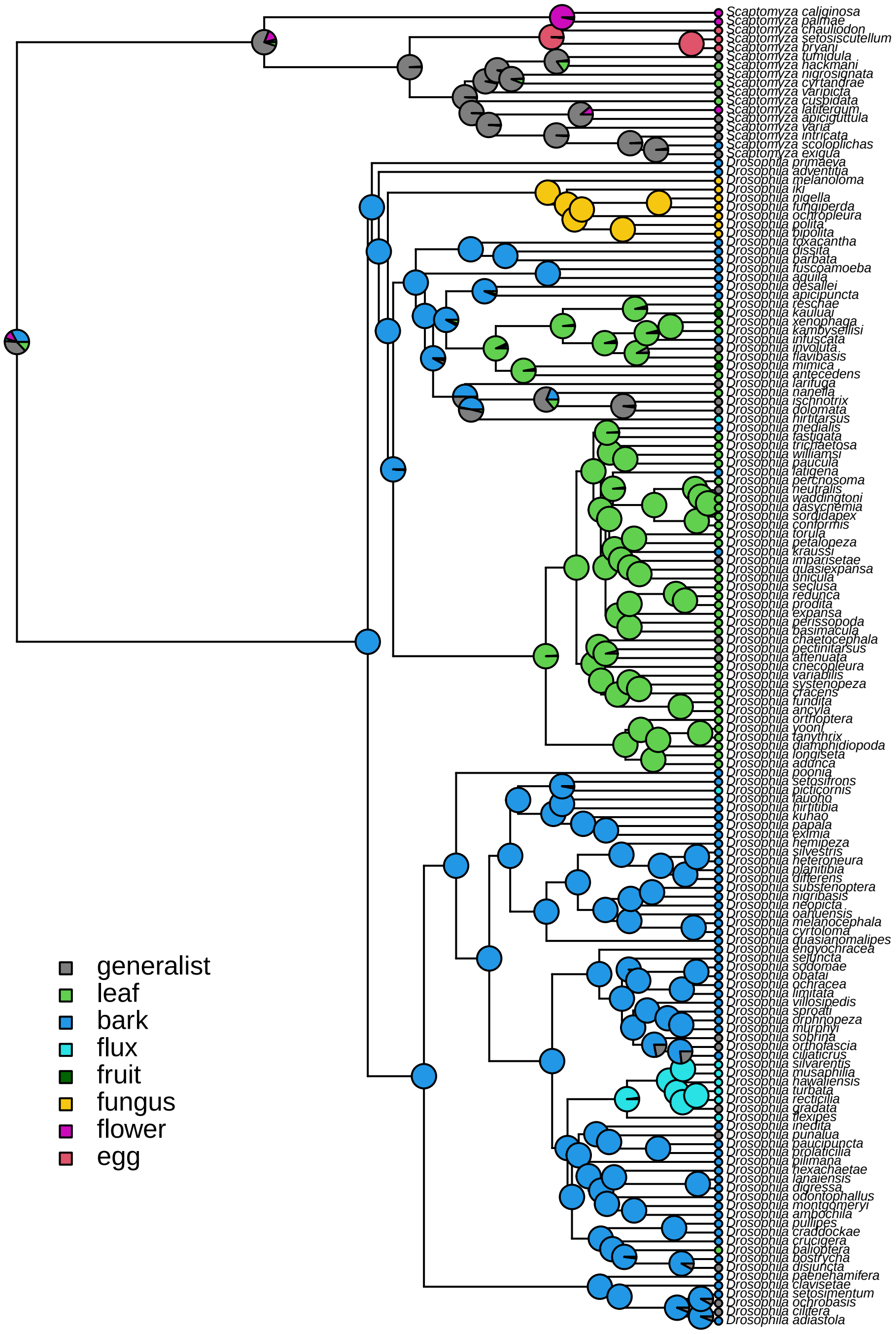

Figure S14: Ancestral state reconstruction of oviposition substrate based on rearing records[9] using stochastic character mapping. Generalist species are defined as those with any two substrates that each comprise >1/4 of rearing records, or any species without one substrate comprising more than >2/3 of rearing records[9]. 'Flux' refers to sap flux breeding, 'egg' refers to spider egg breeding.

### Trait diversification

#### Body length, wing length, and thorax length

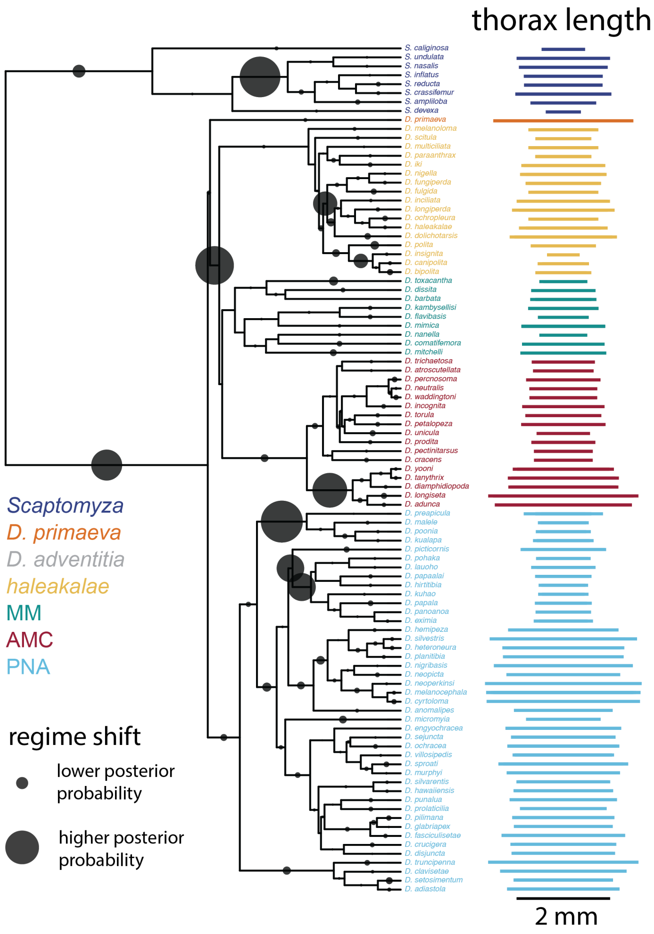

Figure S15: Model of the evolution of thorax length (mm). Data digitized from 26 publications[10–35]. Probable shifts in evolutionary regimes shown by gray circles. Larger circles indicates greater posterior probability that a shift occured on that branch. Distribution of thorax length measurements shown next to tips.

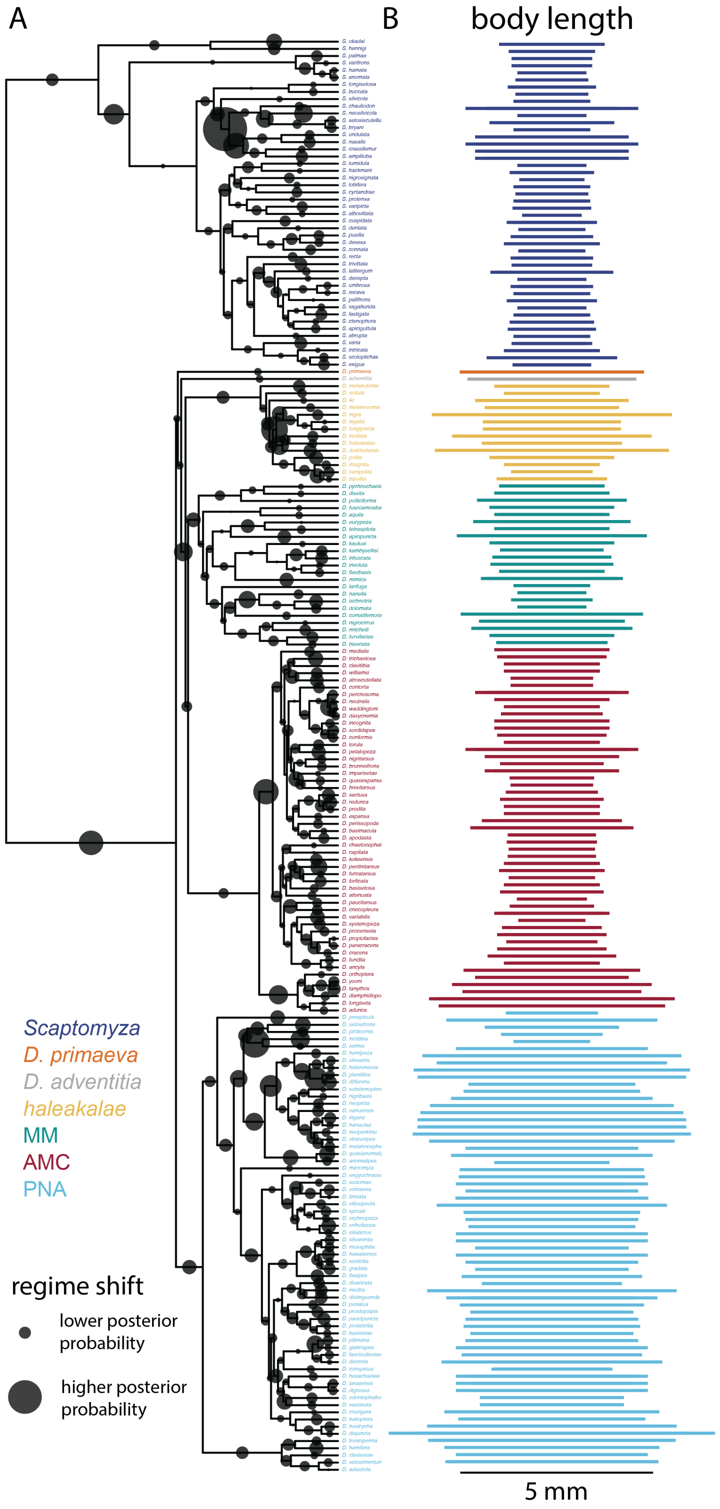

Figure S16: Model of the evolution of body length (mm). Data digitized from 26 publications[10–35]. Probable shifts in evolutionary regimes shown by gray circles. Larger circles indicates greater posterior probability that a shift occured on that branch. Distribution of body length measurements shown next to tips.

### Egg size and shape

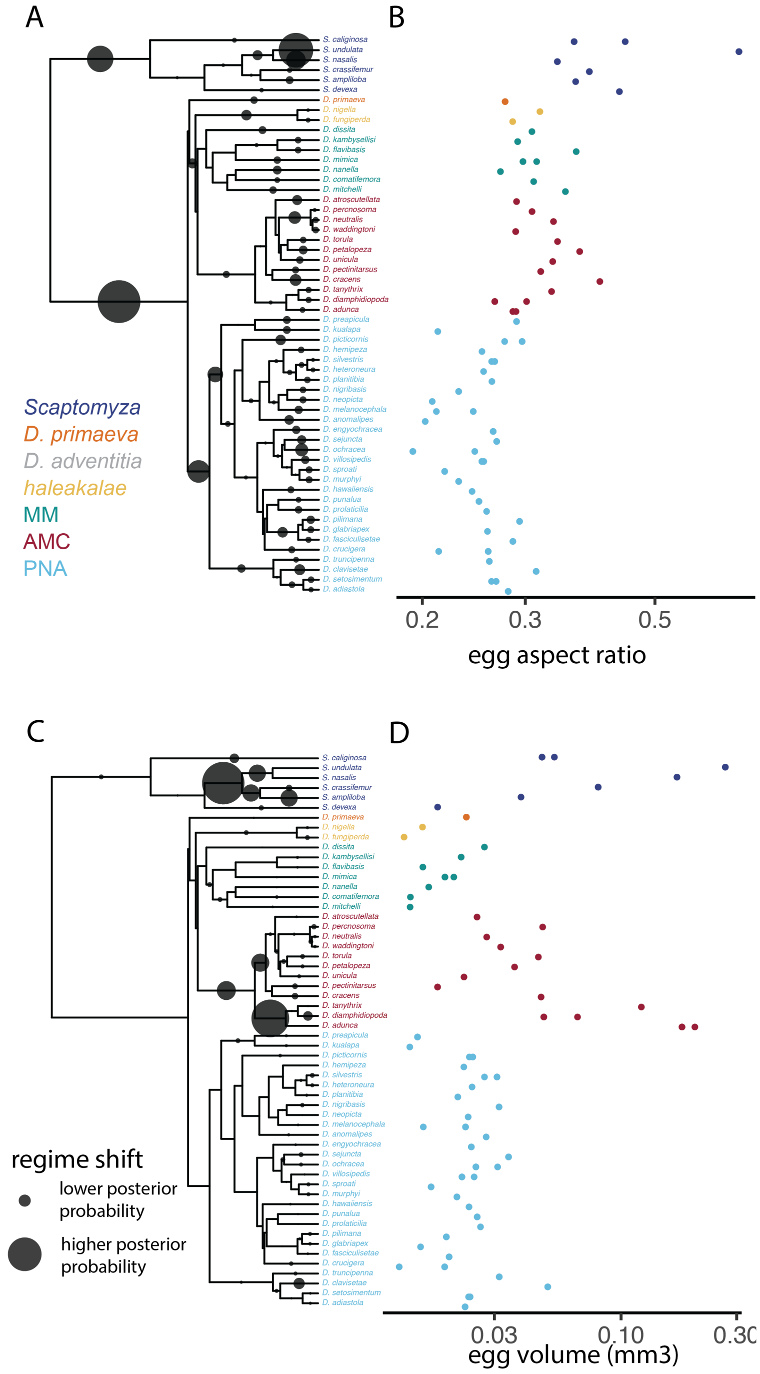

Figure S17: A and C, model of the evolution of egg volume (mm3) and aspect ratio (unitless), probable shifts in evolutionary regimes shown by gray circles. Data digitized from three publications[26,31,35]. Larger circles indicates greater posterior probability that a shift occured on that branch. B and D, Egg volume (mm3) and aspect ratio (unitless), log10 transformed.

### Ovariole number

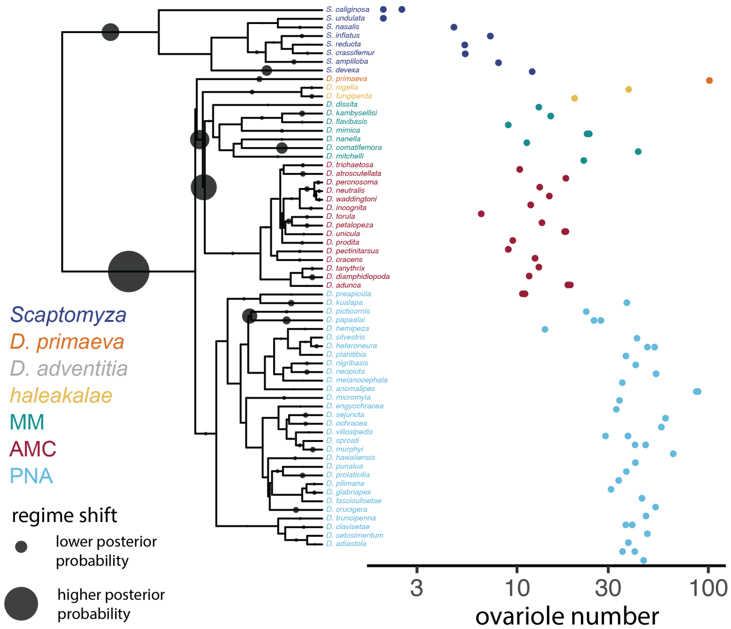

Figure S18: Model of the evolution of ovariole number, probable shifts in evolutionary regimes shown by gray circles. Data digitized from three publications[26,31,35]. Larger circles indicates greater posterior probability that a shift occured on that branch. Ovariole number, log10 transformed, shown adjacent to tips.

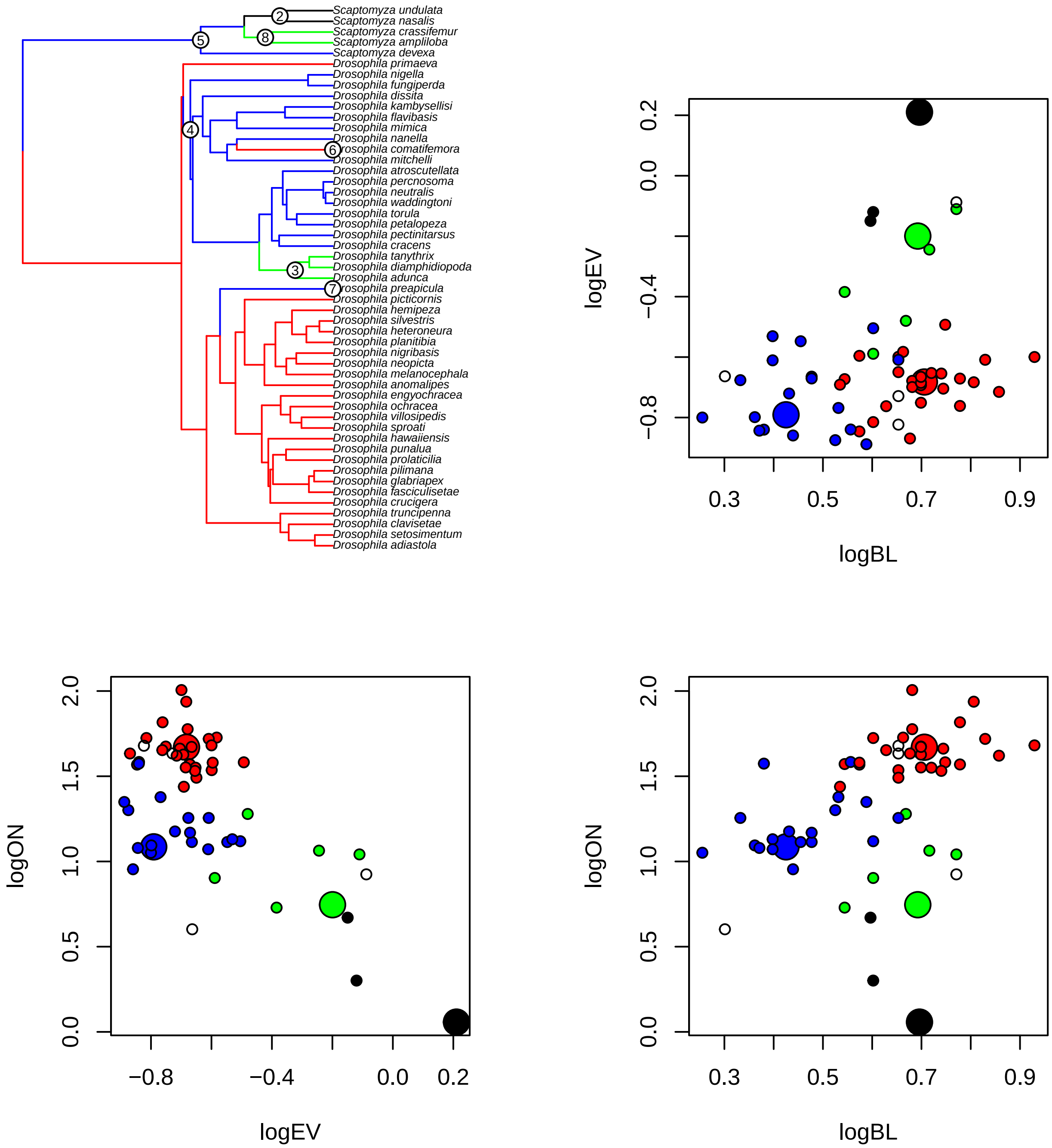

Figure S19: SURFACE estimate of convergent regime shifts in three traits. SURFACE estimates shifts in evolutionary regimes for multiple traits at once, and then assesses whether independent shifts can be combined into convergent regimes. Considering three traits (BL - body length, EV - egg volume, and ON - ovariole number, all log10 transformed, SURFACE finds evidence for eight shifts between four regimes. These can be described as a regime with high EV and low ON, seen in *Scaptomyza* species, a regime with high EV, high BL, and medium ON, seen in *Scaptomyza* and *antopocerus* species, a regime with high ON and high BL, seen in PNA species, *D. primaeva*, and *D. crassifemur*, and a regime with low BL and EV, seen in all others.

**Voucher specimen photos**

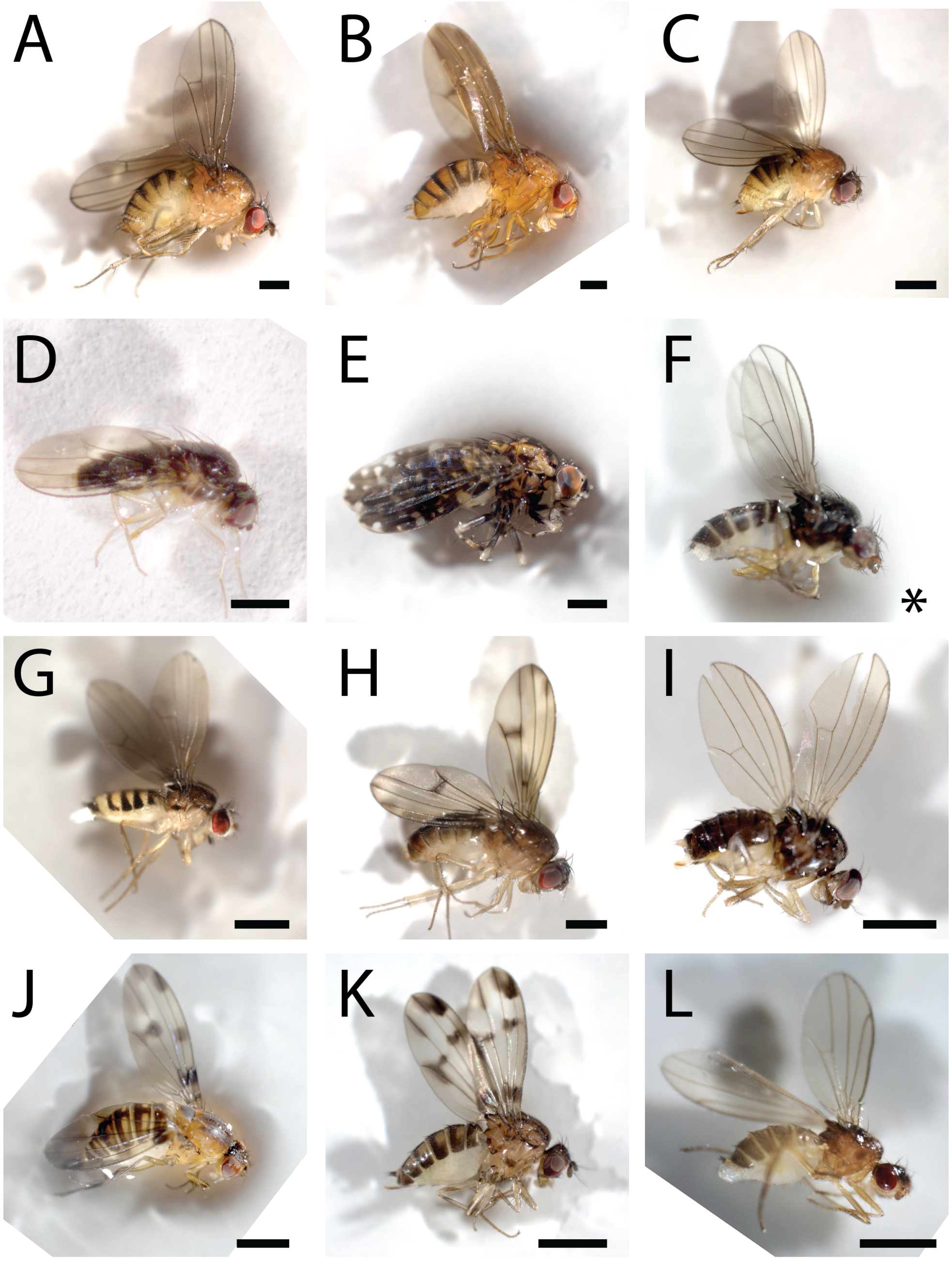

Figure S20: Photos of specimens used for transcriptome sequencing. Species are A, *D. tanythrix*, B, *D. primaeva*, C, *D. atroscutellata*, D, *D.* cf *dives*, E, *D. picticornis*, F, *S. varia*, G, *S. albovittata*, H, *D mimica*, I, *D. nanella*, J, *D. sproati*, K, *D. macrothrix*, L, *S. cyrtandrae*. Scale bar = 1 mm. Asterisk in panel F indicates scale bar failed to be recorded at the time image was captured.

### Supplementary Tables

Table S1: Field collection information for transcriptome sequenced specimens.

| ID | species | general site | locality | collection date | collection method | permit | GPS |
|---|---|---|---|---|---|---|---|
| 16.1-1 | *Drosophila cf dives* | Hawai'i Volcanoes National Park | Bird park | 5/9/2016 | baits | DOFAW I1012; HAVO-2017-SCI-0017 | N19° 26.3512' W155° 18.2225' |
| 040C | *Drosophila mimica* | Hawai'i Volcanoes National Park | Bird park | 4/17/2017 | sweeping Sapindus saponaria leaves | DOFAW I1012; HAVO-2017-SCI-0017 | N19° 26.3512' W155° 18.2225' |
| 055A | *Drosophila macrothrix* | Hawai'i Volcanoes National Park | Ola'a tract, pole 44 | 4/17/2017 | baits | DOFAW I1012; HAVO-2017-SCI-0017 | N19° 27.722' W155° 14.875' |
| 043D | *Drosophila tanythrix* | Hawai'i Volcanoes National Park | Ola'a tract, pole 44 | 4/18/2017 | baits | DOFAW I1012; HAVO-2017-SCI-0017 | N19° 27.722' W155° 14.875' |
| 106A | *Drosophila sproati* | Hawai'i Volcanoes National Park | Ola'a tract, pole 44 | 5/29/2017 | baits | DOFAW I1012; HAVO-2017-SCI-0017 | N19° 27.722' W155° 14.875' |
| 025A | *Drosophila picticornis* | Koke'e State Park | Awa'awapuhi trail | 4/15/2017 | baits | DOFAW I1012; Koke'e state park K2017-2015; I1012; NARS special use; Kaua'i island forest reserves KPI-2017-114 | N22° 08.481' W159° 38.926' |
| 002D | *Drosophila nanella* | Koke'e State Park | Drosophila ditch | 4/13/2017 | sweeping Paisonia Leaves | DOFAW I1012; Koke'e state park K2017-2015; I1012; NARS special use; Kaua'i island forest reserves KPI-2017-114 | N22° 04.795' W159° 40.448' |
| 029A | *Drosophila atroscutellata* | Koke'e State Park | Nualolo trail | 4/16/2017 | sweeping Corynocarpus sp leaves | DOFAW I1012; Koke'e state park K2017-2015; I1012; NARS special use; Kaua'i island forest reserves KPI-2017-114 | N22° 07.801' W159° 39.617' |
| 020A | *Scaptomya varipicta* | Koke'e State Park | Nualolo trail | 4/15/2017 | sweeping Cheirodendron sp. leaves | DOFAW I1012; Koke'e state park K2017-2015; I1012; NARS special use; Kaua'i island forest reserves KPI-2017-114 | N22° 07.801' W159° 39.617' |
| 008D | *Drosophila primaeva* | Koke'e State Park | Pihea trail | 4/14/2017 | baits | DOFAW I1012; Koke'e state park K2017-2015; I1012; NARS special use; Kaua'i island forest reserves KPI-2017-114 | N22° 08.799' W159° 37.074' |
| CFB | *Scaptomya varia* | Koke'e State Park | Pihea trail | 4/14/2017 | collected rotting Clermontia sp flowers | DOFAW I1012; Koke'e state park K2017-2015; I1012; NARS special use; Kaua'i island forest reserves KPI-2017-114 | N22° 08.799' W159° 37.074' |
| 088B | *Scaptomya cyrtandrae* | Stainback Highway | Army road - west | 5/29/2017 | on Cyrtandra platyphylla | DOFAW I1012; NARS special use; Hawai'i island forest reserve access permit | N19° 33.615' W155° 15.010' |

Table S2: DNA barcoding for identification of females.

| individual | sample | species match | reference male | external reference sequence | barcode sequence used for final identification | notes |
|---|---|---|---|---|---|---|
| 029A | 029Atxt | *D. atroscutellata* | yes | yes | COII | |
| 088B | 088Bb | *S. cyrtandrae* | none | yes | COII | |
| 040C | 040Ctxt | *D. mimica* | yes | yes | 16S | |
| 002D | 002Dtxt | *D. nanella* | yes | yes | 16S, COII | |
| 106A | 106Atxt | *D. sproati* | none | none | COII | matched to other females, morphology is distinctive for females in this species |
| 043D | 043Dtxt | *D.tanythrix* | yes | yes | COII | barcode sequences for D. cognata and D. yooni males suggest hidden complexity in this group |
| CFB | CFBb | *S. varia* | yes | yes | COI | |
| 020A | 020Atxt | *S. varipicta* | yes | yes | COII | |
| 16.1-1 | 16.1.4 | *D. cf dives* | none | none | 16S, COII | found no matching reference sequence and no males were caught |

Table S3: Sequencing read counts

| species | individual | sample | tissue | Reads - June 2018 | Reads - July 25, 2018 | Reads - Nov 27, 2018 | Total reads |
|---|---|---|---|---|---|---|---|
| *S. varia* | CFB | CFBb | carcass | | 6,672,052 | | 6,672,052 |
| *S. varia* | CFB | CFBn | head | | 6,311,203 | | 6,311,203 |
| *S. varia* | CFB | CFBo | ovary | | 12,672,693 | | 12,672,693 |
| *S. cyrtandrae* | 088B | 088Bb | carcass | | 9,166,453 | | 9,166,453 |
| *S. cyrtandrae* | 088B | 088Bn | head | | 8,796,864 | | 8,796,864 |
| *S. cyrtandrae* | 088B | 088Bo | ovary | | 9,763,204 | | 9,763,204 |
| *D. sproati* | 106A | 106Atxt | whole fly | 7,432,370 | 6,026,569 | 10,237,744 | 23,696,683 |
| *D. atroscutellata* | 029A | 029Atxt | whole fly | 12,520,922 | 7,801,177 | 19,925,042 | 40,247,141 |
| *D. macrothrix* | 055A | 055Atxt | whole fly | 10,336,762 | 10,313,394 | 19,643,740 | 40,293,896 |
| *D. mimica* | 040C | 040Ctxt | whole fly | 9,471,290 | 8,887,955 | 15,751,511 | 34,110,756 |
| *D. nanella* | 002D | 002Dtxt | whole fly | 10,833,205 | 7,350,705 | 17,211,801 | 35,395,711 |
| *D. picticornis* | 025A | 025Atxt | whole fly | 10,085,602 | 10,177,523 | 16,172,455 | 36,435,580 |
| *D. primaeva* | 008D | 008Dtxt | whole fly | 9,129,075 | 7,583,577 | 13,937,132 | 30,649,784 |
| *D. tanythrix* | 043D | 043Dtxt | whole fly | 11,293,054 | 8,833,878 | 15,667,639 | 35,794,571 |
| *S. varipicta* | 020A | 020Atxt | whole fly | 7,690,349 | 8,004,757 | 14,380,332 | 30,075,438 |
| *D. cf dives* | 16.1-1 | 16.1.1 | ovary | 16,808,131 | | | 16,808,131 |
| *D. cf dives* | 16.1-1 | 16.1.2 | head | 18,215,227 | | | 18,215,227 |
| *D. cf dives* | 16.1-1 | 16.1.4 | body | | 8,773,601 | | 8,773,601 |

Table S4: Genome source information[36,37]

| species | publication | genome source |
|---|---|---|
| *D. virilis* | FlyBase | flybase.org - file dvir-all-transcript-r1.07.fasta |
| *D. grimshawi* | FlyBase | flybase.org - file dgri-all-transcript-r1.05.fasta |
| *D. melanogaster* | FlyBase | flybase.org - file dmel-all-transcript-r6.29.fasta |
| *D. willistoni* | FlyBase | flybase.org - file dwil-all-transcript-r1.05.fasta |
| *D. mojavensis* | FlyBase | flybase.org - file dmoj-all-transcript-r1.04.fasta |
| *D. pseudoobscura* | FlyBase | flybase.org - file dpse-all-transcript-r3.04.fasta |
| *D. annanassae* | FlyBase | flybase.org - file dana-all-transcript-r1.06.fasta |
| *D. murphyi* | Kim et al, 2020 | https://web.stanford.edu/~bkim331/files/genomes/ - accessed January 2021 |
| *S. pallida* | Kim et al, 2020 | https://web.stanford.edu/~bkim331/files/genomes/ - accessed January 2021 |
| *S. hsui* | Kim et al, 2020 | https://web.stanford.edu/~bkim331/files/genomes/ - accessed January 2021 |
| *S. graminum* | Kim et al, 2020 | https://web.stanford.edu/~bkim331/files/genomes/ - accessed January 2021 |
| *S. montana* | Kim et al, 2020 | https://web.stanford.edu/~bkim331/files/genomes/ - accessed January 2021 |

Table S5: Oviposition substrate categories described in rearing records[9]

| substrate category | original substrate listed |
|---|---|
| leaf | leaf |
| leaf | leaf axil |
| leaf | leaf base |
| leaf | live leaf |
| leaf | petiole |
| leaf | frond |
| bark | rachis |
| bark | bark |
| bark | stem |
| bark | wood |
| bark | root |
| bark | branch |
| bark | shoot |
| misc | frass |
| fungus | fungus |
| fruit | fruit |
| fruit | pod |
| egg | spider egg |
| flux | flux |
| flux | soil |
| flower | flower |

Table S6: Effective size of bayou analyses on trait regimes.

| trait | variable | effective size |
|---|---|---|
| body length | k | 791.4 |
| body length | lnL | 401.9 |
| egg aspect ratio | k | 1302.1 |
| egg aspect ratio | lnL | 78.8 |
| egg volume | k | 1378.5 |
| egg volume | lnL | 89.6 |
| ovariole number | k | 1312.5 |
| ovariole number | lnL | 277.0 |
| thorax length | k | 1399.5 |
| thorax length | lnL | 37.6 |
| wing length | k | 378.4 |
| wing length | lnL | 221.9 |
